# Recruitment of Mre11 to recombination sites during meiosis

**DOI:** 10.1101/2025.07.08.663703

**Authors:** Priyanka Priyadarshini, Mahesh Survi, Wael El Yazidi Mouloud, Regina Bohn, Steven Ballet, Neil Hunter, Alexander N. Volkov, Corentin Claeys Bouuaert

**Affiliations:** Louvain Institute of Biomolecular Science and Technology, Université catholique de Louvain, 1348 Louvain-La-Neuve, Belgium; Research Group of Organic Chemistry, Vrije Universiteit Brussel (VUB), Pleinlaan 2, 1050 Brussels, Belgium; Jean Jeener NMR Centre, Vrije Universiteit Brussel (VUB), Pleinlaan 2, 1050 Brussels, Belgium; Howard Hughes Medical Institute, University of California Davis, Davis, CA 95616, USA; Department of Microbiology & Molecular Genetics, University of California Davis, Davis, CA 95616, USA; VIB-VUB Center for Structural Biology, VIB, Pleinlaan 2, 1050 Brussels, Belgium

## Abstract

The Mre11 nuclease, part of the conserved MRX complex involved in the repair of DNA double-strand breaks (DSBs), is also essential to initiate meiotic recombination in budding yeast by promoting Spo11-induced DSBs. Recruitment of Mre11 to meiotic DSB sites depends on Rec114-Mei4 and Mer2 (RMM) that organize the meiotic DSB machinery by a mechanism involving biomolecular condensation. Here, we explored the role of Mre11 during meiosis and its relationship to RMM condensation. We show that both Mre11 and MRX complexes form DNA-dependent, hexanediol sensitive condensates *in vitro. In vivo*, Mre11 assembles into DNA damage-dependent foci in vegetative cells and DSB-independent foci in meiotic cells. *In vitro* condensates and *in vivo* foci both depend on the C-terminal intrinsically-disordered region (IDR) of Mre11. Importantly, while the Mre11 IDR is dispensable for vegetative DNA repair it is essential during meiosis. The C-terminus of Mre11 forms a short α-helix that binds a conserved region of Mer2, and mutating residues within this interface reduces Mre11 foci and DSB formation. Finally, we identified a SUMO-interacting motif within the Mre11 IDR that enhances recruitment of Mre11 during meiosis and facilitates DSB formation. This work identifies multiple mechanisms that collaborate to recruit Mre11 during meiosis to initiate recombination.

## Introduction

Cells continuously experience exogenous and endogenous stress leading to cytotoxic lesions such as DNA double-strand breaks (DSBs)^1,2^. If not repaired promptly and accurately, DSBs can compromise genome stability resulting in cellular dysfunction, tumorigenesis, or cell death^3^. While DSBs are generally considered detrimental for genome integrity, several instances of programmed DSBs that serve a physiological purpose are also well-documented, including V(D)J recombination during lymphocyte development, immunoglobulin class-switching, and meiotic recombination^4^.

During meiosis, recombination initiated by programmed DNA DSBs serves two critical functions: it ensures accurate segregation of genetic material by forming physical connections between homologous chromosomes and increases genetic diversity through allelic shuffling^5,6^. Meiotic DSBs are generated by the highly conserved type II topoisomerase-like protein Spo11 that acts in a consortium with several accessory factors, referred to as DSB proteins^7,8^.

In *Saccharomyces cerevisiae*, the nine known DSB proteins include Rec114-Mei4 and Mer2. These RMM proteins are proposed to assemble into nucleoprotein condensates along meiotic chromosomes^9^ that act as sub-cellular compartments that recruit other DSB proteins to stimulate DNA cleavage by controlling Spo11 dimerization^10–12^. This condensation model provides a mechanism that ties DSB formation to the loop-axis organization of meiotic chromosomes^13,14^.

In *S. cerevisiae*, DSB formation also depends on the Mre11-Rad50-Xrs2 (MRX) complex^15–18^. In addition, MRX initiates the resection of DSB ends in both meiotic and mitotically cycling cells, through the endonuclease and exonuclease activities of Mre11^19–25^. Further, the MRX complex has notable roles in telomere maintenance, stabilization of replication forks, and viral infection^26–28^.

While the roles of MRX in genome stability have been studied extensively, its role in meiotic DSB formation remains poorly understood. Here, we show that Mre11 forms dynamic condensates *in vitro* dependent on its C-terminal disordered tail. The Mre11 C-terminus was previously shown to be required for meiotic DSB formation^29,30^ and was recently found to bind Spo11^31^. We demonstrate that the C-terminus of Mre11 also binds directly to Mer2 and contains a SUMO-interacting motif that also facilitates its recruitment to DSB sites. Thus, our work delineates multiple mechanisms whereby the Mre11 C-terminal tail specifically promotes meiotic DSB formation.

## Results

### Mre11 and the MRX complex assemble DNA-dependent condensates in vitro

During meiosis and in mitotically cycling cells exposed to DNA damage, budding yeast Mre11 forms numerous foci visible by immunofluorescence microscopy^30,32,33^. *In vitro*, Mre11-Rad50 complexes have been shown to form higher-order assemblies on DNA, mediated by Rad50 oligomerization^34^. However, whether Mre11 itself participates in the assembly of higher-order structures has not been established.

To investigate the biochemical properties of Mre11 and the MRX complex, we purified Alexa488-labeled Mre11 and MRX complexes with an eGFP tag at the N-terminus of Mre11 (**Figure S1A and S2A**) and asked whether they undergo condensation *in vitro*. In the presence of plasmid DNA, we found that Mre11 and MRX form large clusters visible by fluorescence microscopy (**Figures 1A and 1C**). While foci were almost absent when DNA was omitted, low concentrations of DNA led to few foci of very high intensity, suggesting that the limited number of DNA molecules provide nucleation points for Mre11 to agglomerate (**Figure S1B**). Time-course analysis showed a progressive increase in foci number and intensity over time (**Figure S1C**); and protein titration revealed a steep increase in foci intensity above 200 nM Mre11 (**Figure S1D**). A similar concentration-dependent increase in foci intensity was also observed for MRX condensates (**Figure S2B**). Mre11 and MRX condensation was also promoted in the presence of crowding agents (**Figures 1E and 1F**). Mre11 condensation was enhanced in the presence of divalent metal ions, likely through charge neutralization (**Figure S1E**). To address whether the condensates are reversible, we assembled condensates for 20 minutes and treated the reactions with DNase I. While nuclease treatment strongly reduced the numbers of Mre11 and MRX foci, the minority of Mre11 foci that resisted nuclease treatment tended to accumulate more protein (**Figures 1B and 1D**). This observation is consistent with the idea that, following initial nucleation by DNA, Mre11 and MRX condensates grow primarily through protein-protein interactions.

**Figure 1:**
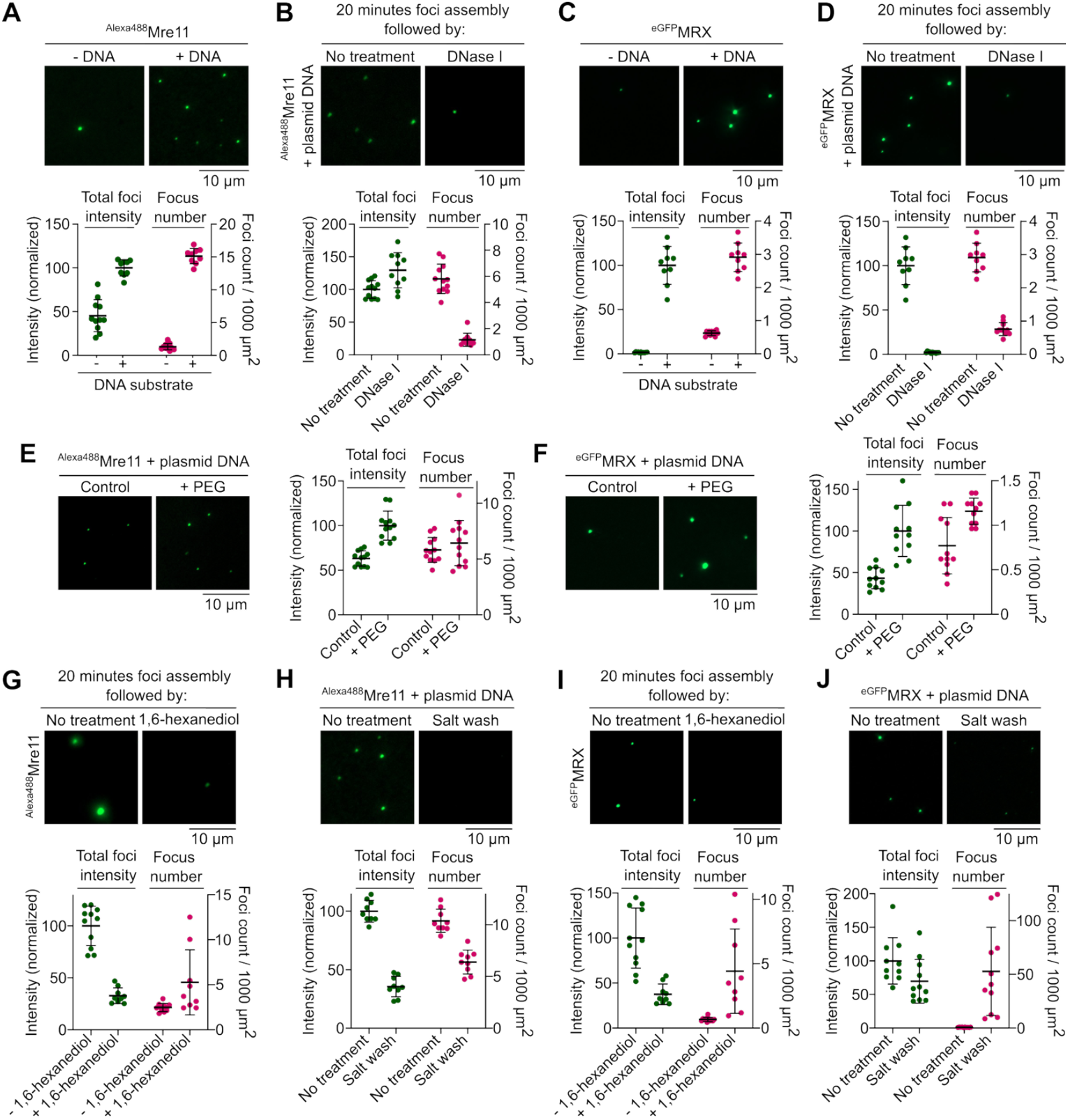
Mre11 and the MRX complex assemble DNA-dependent condensates *in vitro*. **(A, C)** Effect of the presence of plasmid DNA (pUC19) on condensation of (A) Alexa488-labelled Mre11 and (C) eGFP-tagged MRX. **(B, D)** Effect of DNase I on (B) Mre11 and (D) MRX condensates. Condensates were pre-assembled for 20 minutes with 5.7 nM pUC19 prior to challenge. Quantification represents total foci intensity (green) in a field of view and total number of foci per 1000 μm^2^ (magenta). Foci intensity is normalized to the mean intensity of samples with DNA for A, C and untreated samples for B, D. Error bars represent mean ± SD from 8-12 fields of view. **(E, F)** Effect of the presence of 5% PEG-3350 on (E) Mre11 and (F) MRX condensate formation. Quantification represents total foci intensity (green) in a field of view (normalized to the mean of +PEG images) and total number of foci per 1000 μm^2^ (magenta). Error bars represent mean ± SD from 10-12 fields of view. **(G, I)** Effect of challenge with 5% 1,6-hexanediol and **(H, J)** 0.5 M NaCl on (G, H) Mre11 and (H, J) MRX condensates. Condensates were assembled for 20 minutes prior to challenge and incubated for 15 minutes and 10 minutes, respectively, after challenge. In the control reaction, an equivalent volume (1/20) of water was added instead of hexanediol. Quantification shows foci intensity (normalized to the mean of untreated samples) and total number of foci per 1000 μm^2^ (magenta). Error bars represent mean ± SD from 10-12 fields of view.

To further test this interpretation, we asked whether Mre11 and MRX condensates are sensitive to 1,6-hexanediol, an aliphatic alcohol that often dissolves biomolecular condensates^35^. Ten-minute treatment with 5% hexanediol led to a strong reduction in foci intensity (**Figures 1G and 1I**). Similarly, brief exposure to 500 mM NaCl strongly reduced foci intensity (**Figures 1H and 1J**). Hence, Mre11 and MRX nucleoprotein condensates are stabilized through a combination of ionic and weak hydrophobic interactions.

### The C-terminal IDR of Mre11 is required for condensation

Mre11 consists of an N-terminal phosphodiesterase domain, a capping domain followed by a Rad50 binding domain, and a C-terminal tail, predicted to be largely disordered (**Figures 2A and 2B**)^36–38^. The last 15 residues form a short, conserved helix. It was previously shown that the C-terminal 49 amino acids are dispensable for mitotic DSB repair but important for meiotic DSB formation^29^.

**Figure 2:**
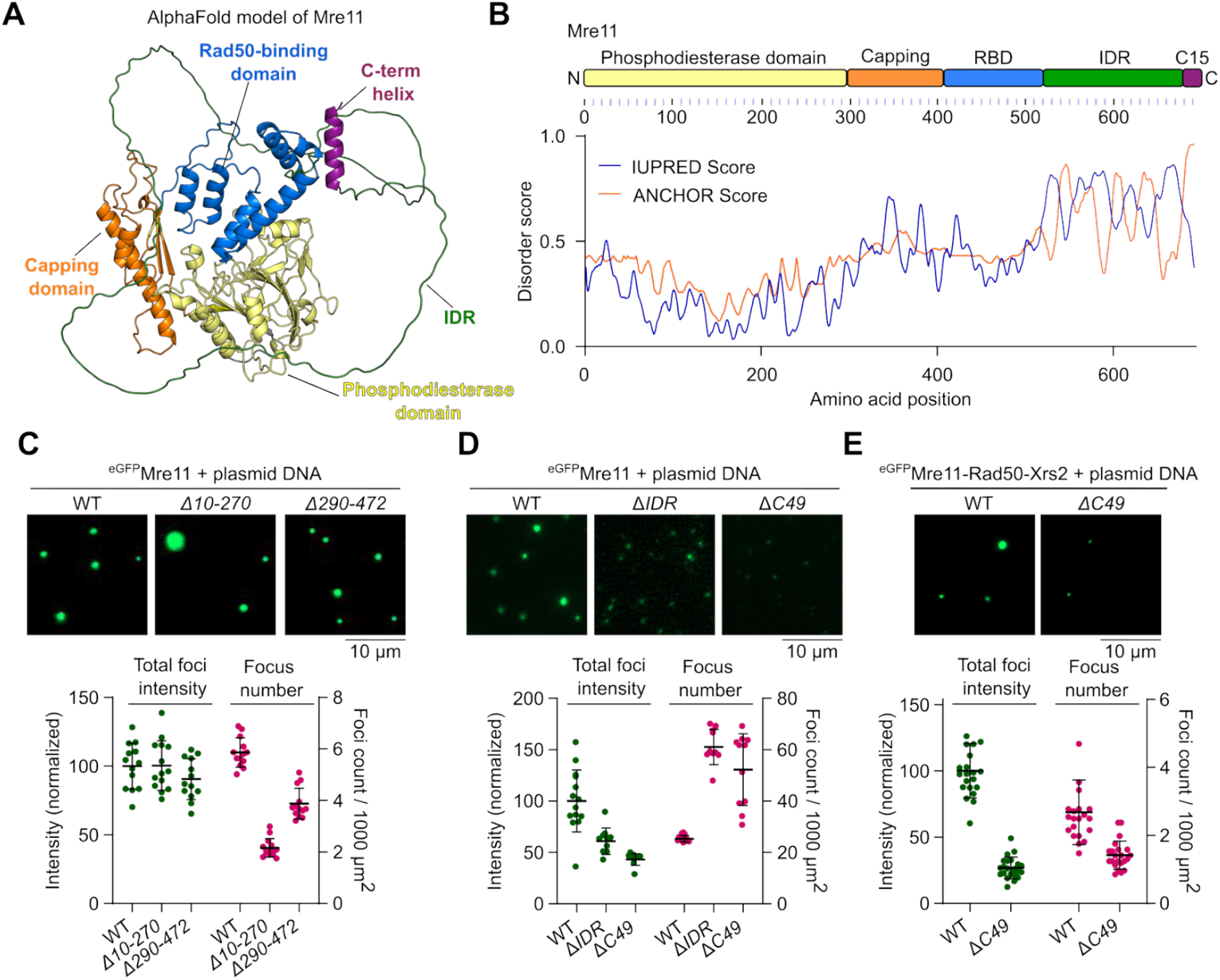
The C-terminal IDR of Mre11 is required for condensation. **(A)** AlphaFold2 predicted model of Mre11 (AF-P32829-F1-v4). **(B)** Domain structure of Mre11 and protein disorder prediction using IUPred server^73^. The ANCHOR score predicts the transition probability from a disordered to an ordered structure dependent on a binding partner. **(C)** Effect of deleting most of the phosphodiesterase domain (*Δ10-270*), or the capping domain and Rad50-binding domain (*Δ290-472*), **(D)** deleting the disordered region (*ΔIDR*, residues 524-677) and C-terminus (*ΔC49*, residues 644-692) on Mre11 condensation, and (**E**) deleting the C-terminus (*ΔC49*) on MRX condensation. Reactions in (C) and (D) contained 200 nM Mre11. Quantification represents total fluorescence intensity (green) in a field of view and total number of foci per 1000 μm^2^ (magenta). Foci intensity is normalized to the mean foci intensity of wild type Mre11 or MRX. Error bars represent mean ± SD from 10-14 fields of view for (C) and (D), and 21 fields of view for (E).

To identify the domain(s) required for Mre11 condensation, we purified eGFP-tagged truncations (**Figures S3A and S3B**). Deletion of residues 10-270 that removes most of the phosphodiesterase domain and residues 290-472 that removes the capping domain and most of the Rad50-binding domain led to reduced numbers of foci that retained high intensity (**Figure 2C)**. In contrast, deleting the disordered region (residues 524-677, Δ*IDR*) or C-terminal 49 residues (residues 644-692, Δ*C49*) strongly reduced Mre11 foci intensity (**Figure 2D)**. Similarly, the Mre11 C-terminal extremity was also required for efficient focus formation of eGFP-tagged MRX complexes (**Figure 2E**).

To understand the effects of Mre11 truncations on DNA-binding activity, we quantified binding to a pUC19 plasmid substrate by gel shift analysis (**Figures S3C and S3D**). All of the truncations retained significant DNA-binding activity. Similar to Mre11^WT^, Mre11^*Δ10-270*^, and Mre11^*Δ290-472*^ produced fast-migrating complexes at concentrations up to ∼25 nM protein, then formed complexes of reduced mobility at higher protein concentration, presumably indicative of higher-order assemblies. However, these species of reduced electrophoretic mobility were absent with Mre11^Δ*IDR*^ and Mre11^Δ*C49*^.

While the Mre11 IDR is required for efficient Mre11 condensation, purified eGFP-tagged Mre11^IDR^ alone failed to form condensates and showed reduced DNA binding as compared to the WT or Mre11^Δ*IDR*^ (**Figures S3E-S3F**). We conclude that, while protein-DNA interactions are important for the formation of Mre11 foci *in vitro*, multivalent protein-protein interaction through low-complexity regions contribute to condensation.

### In mitotically cycling cells, the Mre11 IDR is required for focus formation but dispensable for DNA repair

To investigate the relationship between DNA-damage induced Mre11 foci and DNA repair, we treated vegetative yeast cells expressing myc-tagged Mre11 with methylmethanesulfonate (MMS) and visualized focus formation by immunofluorescence microscopy. A brief exposure to 5% 1,6-hexanediol readily dissolved MMS-induced foci, which instead coalesced into a few aggregates of high intensity (**Figures 3A and 3B)**.

**Figure 3:**
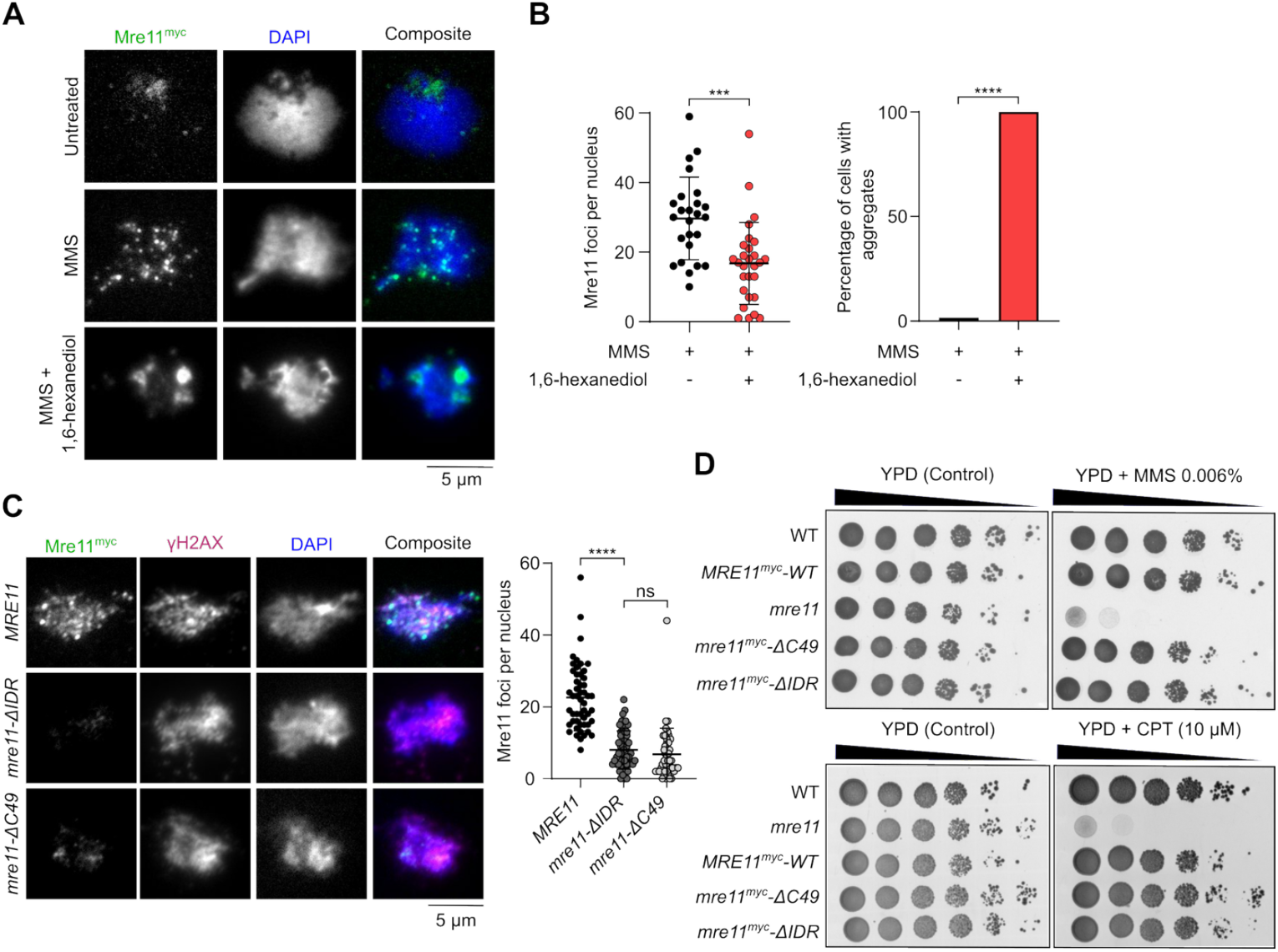
The Mre11 IDR is required for vegetative foci formation but dispensable for DNA repair. **(A)** Effect of 1,6-hexanediol treatment on MMS-induced Mre11^myc^ foci visualized by immunofluorescence analysis of vegetatively growing yeast nuclear spreads **(B)** Quantification of number of Mre11 foci (left) and Mre11 aggregates (right) before and after 1,6-hexanediol treatment. Error bar shows mean ± SD for at least 25 cells per strain. **(C)** Effect of Mre11^myc^ truncations on MMS-induced foci formation as visualized by immunofluorescence. Error bars show mean ± SD of Mre11^myc^ foci for *WT* (n = 51), *mre11-*Δ*IDR* (n = 52), and *mre11-*Δ*C49* (n = 45). **(D)** Sensitivity of wild-type and truncated Mre11^myc^ strains to methylmethanesulfonate (MMS) and camptothecin (CPT). Ten-fold serial dilutions from saturated cultures are shown, with dilutions on YPD plates as control.

Deletion of the IDR (*mre11-ΔIDR*) or C-terminal 49 residues (*mre11-ΔC49*) severely reduced MMS-induced focus formation relative to wild type (**Figure 3C, S4B, S4C**). Despite being important for focus formation, the C-terminus of Mre11 was previously shown to be dispensable for vegetative DNA repair^29^. Consistently, the *mre11-ΔC49* mutant was resistant to treatment with MMS or camptothecin (CPT). Similarly, *mre11-ΔIDR* mutants were as resistant to MMS or CPT as wild-type cells (**Figure 3D**). Hence, the formation of condensate-like Mre11 foci is dispensable for DSB repair in mitotically cycling cells.

### Mer2 condensates directly recruit Mre11 during meiosis

During meiosis, Mre11 appears as discrete foci in immunofluorescence staining of prophase I chromosome spreads^30,31^. Analogous to mitotic cells, a brief exposure to 1,6-hexanediol disassembles these foci (**Figures 4A and S4A**). However, unlike vegetative conditions, we did not detect large Mre11 aggregates upon 1,6-hexanediol treatment during meiosis.

**Figure 4:**
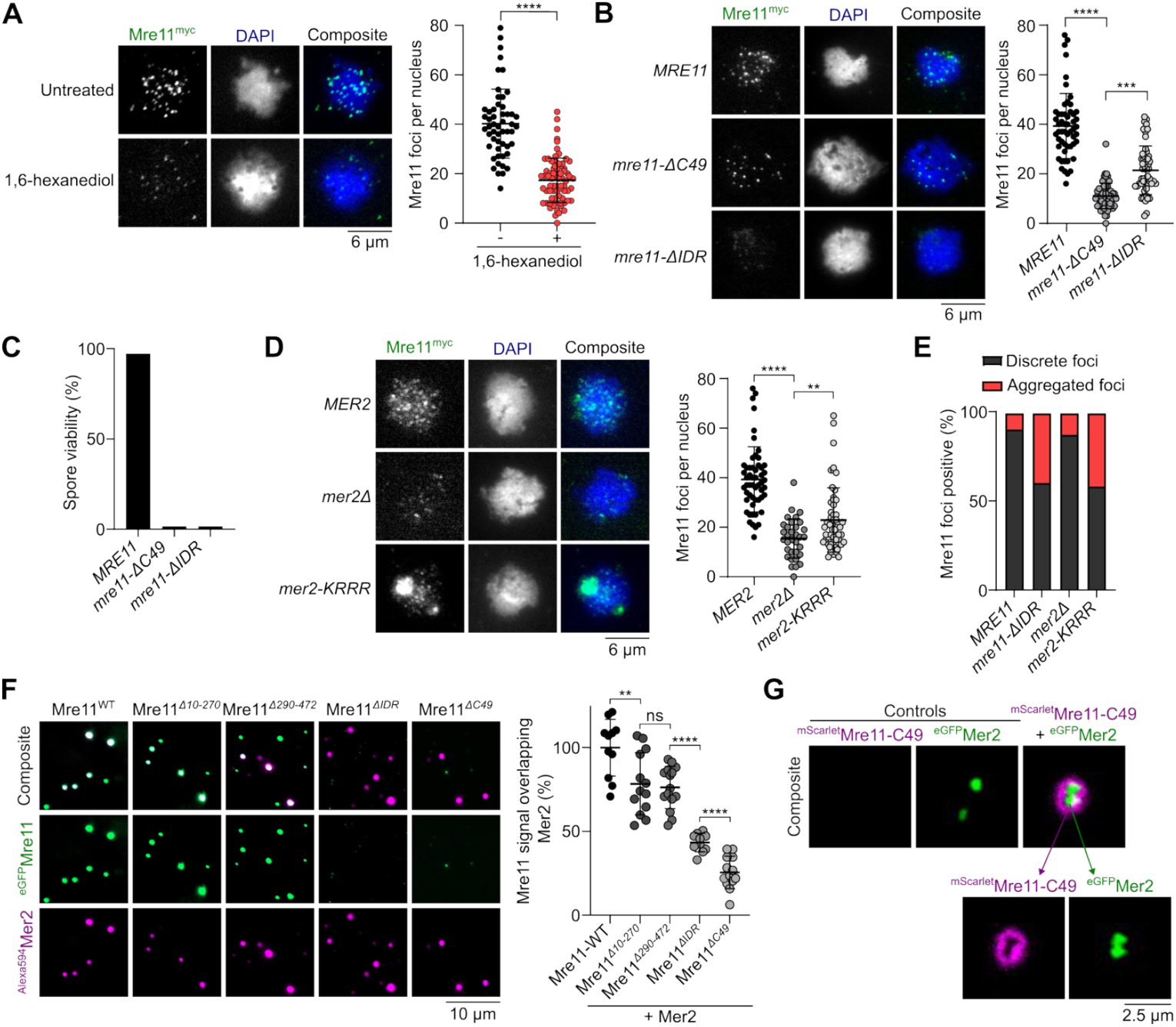
Mer2 recruits Mre11 during meiosis. **(A)** Effect of 1,6-hexanediol treatment on meiotic Mre11^myc^ foci visualized by immunofluorescence analysis of yeast nuclear spreads 4 hours after transfer to SPM. Quantification shows mean and SD for untreated (n = 53) and 1,6-hexanediol treated cells (n = 73). **(B, D)** Immunofluorescence on meiotic nuclear spreads for myc-tagged (B) *MRE11-WT, mre11-ΔC49, mre11-ΔIDR* and (D) *MRE11-WT* in *MER2, mer2Δ*, and *mer2-KRRR* strains. Quantification shows mean and SD of Mre11^myc^ foci for *WT* (n = 55), *mre11-*Δ*C49* (n = 60), *mre11-*Δ*IDR* (n = 53), *MER2* (n = 55), *mer2Δ* (n = 33), and *mer2-KRRR* (n = 57). **(C)** Spore viabilities of strains expressing wild-type or truncated Mre11^myc^. At least 22 tetrads were dissected for each strain (n ≥ 88 spores). (**E**) Quantification of sub-population of cells showing discrete Mre11 foci and aggregated Mre11 foci in indicated strains. Images used for analysis were from the same experiments in (B) and (D). **(F)** Colocalization of fluorescently-labelled Mer2 with wild-type or truncated Mre11. Reactions containing 200 nM of ^Alexa594^Mer2 and ^eGFP^Mre11 were assembled separately for 10 minutes then mixed at 1:1 ratio for 30 minutes prior to imaging. Quantification shows the fraction of Mre11 foci overlapping Mer2 foci (white) in a field of view, normalized to Mre11-WT. Error bars represent mean ± SD from 10-15 fields of view. **(G)** Colocalization of fluorescently-labelled Mre11-C49 and Mer2. Condensates were assembled by mixing 800 nM of ^mScarlet^Mre11-C49 and 200 nM of ^eGFP^Mer2 for 30 minutes prior to imaging. Controls reactions contained the same concentration but lacked either ^eGFP^Mer2 or ^mScarlet^Mre11-C49, respectively.

While Mre11 foci in mitotic cells are formed in response to DNA damage, meiotic Mre11 foci are formed independently of DSBs in cells carrying the catalytically inactive *spo11-Y135F* allele^39^ (**Figures S4D and S4E**). In a wild-type background, the number of Mre11 foci tended to decrease in late meiotic prophase, while foci accumulated in a *spo11-Y135F* mutant (**Figure S4D**). Consistent with the known meiotic function of the Mre11 C-terminus^29^, *mre11-C49* cells produced much fewer foci in meiosis than the wild-type *MRE11* cells (**Figure 4B**). However, the few foci observed had an intensity similar to the wild type, suggesting that the Mre11 C-terminus is important for the nucleation of Mre11 foci, but not their growth. In contrast, the *mre11-ΔIDR* strain produced fewer and much weaker foci. Approximately 40% of the cells also showed accumulation of fluorescent signal in discrete aggregates (**Figure 4B, 4E, S4B, S4C**). As expected, meiosis failed in both *mre11-C49* and *mre11-ΔIDR* strains, producing only dead spores (**Figure 4C**).

Meiotic foci of Mre11 depend on most other DSB proteins, including RMM^40^, and direct interactions have been demonstrated between Mre11, Mer2^41^, and Spo11^31^. To address whether the recruitment of Mre11 during meiosis depends on RMM condensation, we quantified Mre11 foci formation in a DNA-binding defective mutant of Mer2 (*mer2-KRRR*) that compromises Mer2 focus formation *in vitro* and *in vivo* and abolishes DSB formation^9^. Similar to *mer2Δ* strains, the formation of Mre11 foci was strongly reduced in a *mer2-KRRR* mutant (**Figure 4D**), despite no effect on protein levels (**Figures S4F and S4G**). In *mer2Δ* and *mer2-KRRR* backgrounds, Mre11 localization was analogous to the *mre11-ΔIDR* mutant, with few foci of low intensity and accumulation within a large aggregate (**Figures 4D and 4E**).

To test whether Mer2 condensates can directly recruit Mre11 *in vitro*, we used fluorescently labelled Mre11 and Mer2, assembled their respective condensates separately on DNA substrates, and imaged the foci by microscopy. When Mer2 and Mre11 condensates were first assembled separately on a DNA substrate and then incubated together prior to imaging, we observed nearly absolute colocalization between the two proteins, supporting the hypothesis that Mer2 condensates recruit Mre11 through direct protein-protein interactions (**Figure 4F**).

When these experiments were performed with the Mre11-ΔC49 truncation and wild-type Mer2, we observed strongly reduced colocalization as compared to wild-type Mre11 (**Figure 4F**), suggesting that the C-terminus of Mre11 is important for interaction with Mer2. While C-terminus of Mre11 does not form condensates independently, in the presence of Mer2, it forms a halo-like structure around Mer2 condensates (**Figure 4G**). This suggests that the interaction between Mer2 and the Mre11 C-terminus provides a nucleation site that is sufficient for further Mre11 recruitment through self-association.

### The C-terminus of Mre11 binds a conserved motif of Mer2

To delve deeper into the interaction between Mre11 and Mer2, we used AlphaFold2^42^ to model interaction between Mre11-C49 and a Mer2 tetramer (previously defined as the Mer2 oligomeric state^9^)^43^. AlphaFold produced a model in which the terminal α-helix of Mre11 is positioned close to the N-terminal end of the Mer2 coiled coil (**Figure 5A)**. Interestingly, this region contains a conserved sequence motif 1 (SSM1)^44^ that was previously implicated in Mre11 binding^41^. Despite the moderate confidence in the predicted interaction surface, modeling of pairs of Mer2-Mre11 homologs in different species of Saccharomycetaceae produced similar models for two out of three tested species (**Figure S5A**).

**Figure 5:**
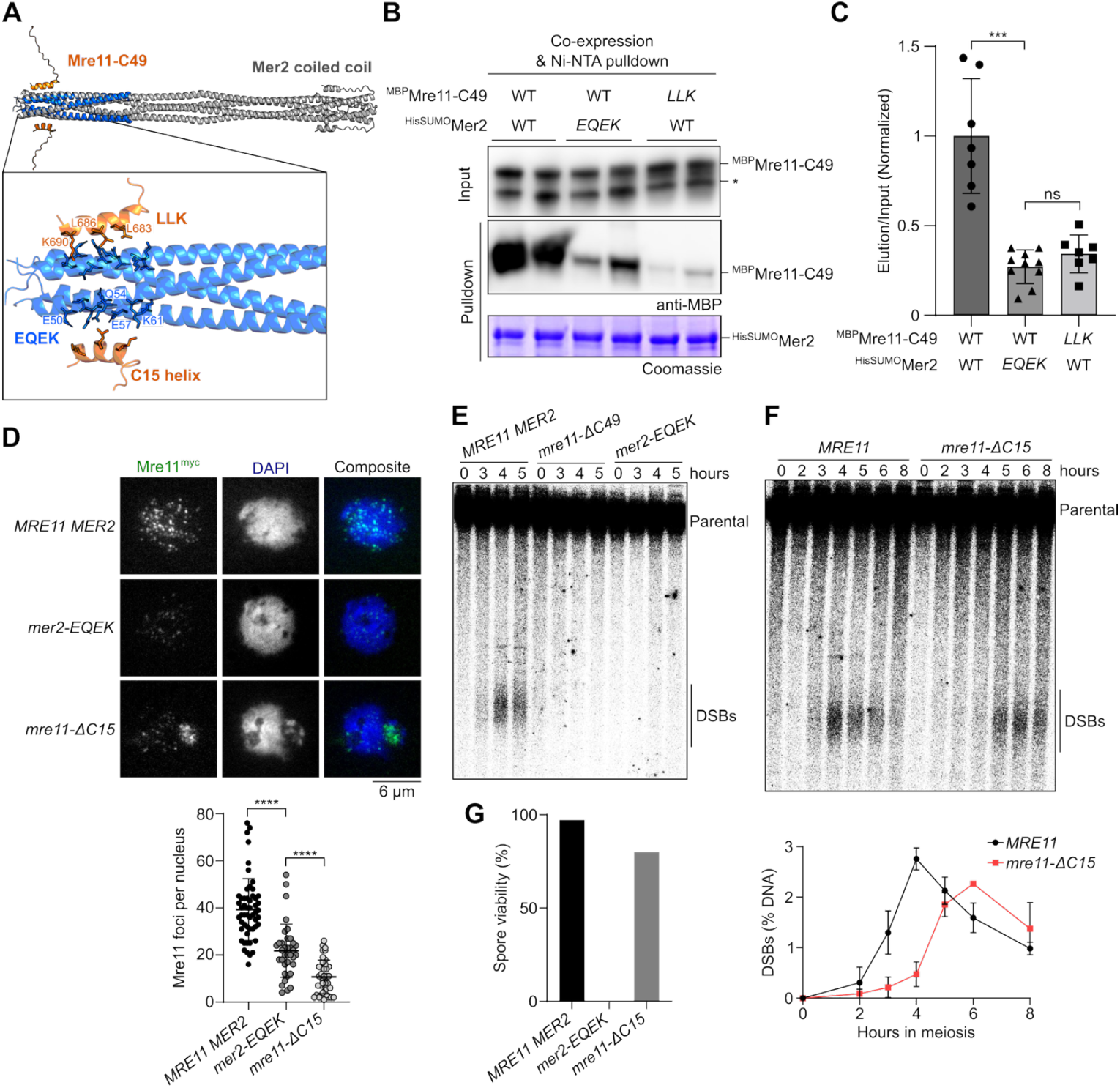
The C-terminus of Mre11 binds a conserved motif of Mer2. **(A)** AlphaFold2 predicted model of Mre11 C-terminal residues (644-692, Mre11-C49) (orange) and Mer2 (41-100) (blue), aligned to the Mer2 coiled-coil domain (residues 41-247) (grey). Magnified-view shows amino acid residues at the predicted interaction surface, Mre11 residues L683, L686, and K690 (LLK) and Mer2 residues E50, Q54, E57, and K61 (EQEK). **(B)** Co-expression based pulldown assay between wild type and mutant ^MBP^Mre11-C49 (prey) and ^HisSUMO^Mer2 (bait). The LLK and EQEK mutants have the respective residues mutated to alanine. Asterisk (*) denotes free MBP. **(C)** Quantification of anti-MBP elution/input signal from pulldown in panel B, normalized to the wild type. Error bars represent mean ± SD from at least seven independent samples. **(D)** Immunofluorescence on meiotic nuclear spreads of Mre11^myc^ in WT, *mer2-EQEK*, and *mre11-*Δ*C15* strains. Quantification of Mre11^myc^ foci in WT (n = 54), *mer2-EQEK* (n = 36), and *mre11-*Δ*C15* (n = 38). **(E, F)** Southern blot analysis of meiotic DSB formation at the *GAT1* hotspot. Quantification of panel F show mean and range from n = 2 experiments. **(G)** Spore viabilities of wild-type and mutant strains (n ≥ 88 spores).

In the AlphaFold model, the Mre11 C-terminal helix binds anti-parallel to the Mer2 coiled coil (**Figure 5A**). Mer2 residues E50, Q54, E57, and K61 (hereby referred to as EQEK) project towards the Mre11 C15 helix, and Mre11 residues L683, L686, and K690 (hereby referred to as LLK) point towards the SSM1 region of Mer2. These residues are well-conserved across Saccharomycetaceae (**Figures S5B and S5D**).

To test this model, we co-expressed MBP-tagged Mre11-C49 peptide and HisSUMO-tagged Mer2 in *E. coli* and quantified protein interactions by NiNTA pulldown. While the interaction was relatively weak, anti-MBP immunoblot analysis confirmed that Mre11-C49 directly binds Mer2 (**Figure 5B**). Consistent with the AlphaFold model, alanine mutations of the Mer2-EQEK or Mre11-LLK residues strongly decreased the interaction between the two partners (**Figures 5B and 5C**).

To establish the relevance of this interaction for Mre11 focus formation *in vivo*, we performed immunofluorescence analysis of the Mer2-EQEK mutant protein. Consistent with our *in vitro* analysis, focus formation of Mer2-EQEK was diminished compared to wild-type Mre11 by approximately 45% (**Figure 5D**). Similarly, deleting the C-terminal α-helix of Mre11 (*mre11-ΔC15*) also reduced focus formation by approximately 73%. (**Figure 5D**). Importantly, reduced focus formation was not due to reduced levels of Mer2-EQEK or Mre11-ΔC15 proteins, which were similar to wild-type Mre11 **(Figures S6A-S6C)**.

To address whether the interaction between Mer2 and Mre11 is important for their meiotic function, we analyzed DSB formation at the *GAT1* hotspot by Southern blotting. Similar to the *mre11-ΔC49* mutant, little or no DSB signal was detected in *mer2-EQEK* cells (**Figure 5E**), while the *mre11-ΔC15* mutation caused reduced and delayed DSB formation (**Figure 5F**), even though the progression of meiosis was unchanged (**Figure S6D**). Consistently, *mer2-EQEK* mutant spores were completely inviable while the *mre11-ΔC15* truncation showed 80% spore viability (**Figures 5G**). Since *mre11-ΔC15* does not phenocopy the *mre11-ΔC49* and *mer2-EQEK* mutations, we conclude that the Mer2 SSM1 motif and the C-terminus of Mre11 exert other meiotic functions than the direct Mer2-Mre11 interaction identified here.

### The Mre11 C-terminus contains a novel SUMO-interaction motif

Several DSB proteins, including Mer2, were shown to be SUMOylated during meiosis, and SUMOylation regulates all aspect of meiotic prophase I, including DSB formation^45^. Notably, SUMOylation sites were mapped in the SSM1 region of Mer2 that also harbors the EQEK residues.

Mre11 has two previously identified SUMO-interacting motifs located in its phosphodiesterase domain (SIM1 and SIM2) that have been implicated in DSB repair in both mitotic and meiotic cells^46^. Using the GPS-SUMO tool^47^, we identified a third potential SIM, here called SIM3 (IIMVS), located towards the end of the Mre11 IDR (**Figures 6A and S7A**). AlphaFold prediction yielded a high-confidence model in which Mre11-SIM3, which is otherwise disordered, assumes a β-sheet structure in the presence of Smt3, as expected from a *bona fide* SIM^48^ (**Figures S7B-S7D**).

**Figure 6:**
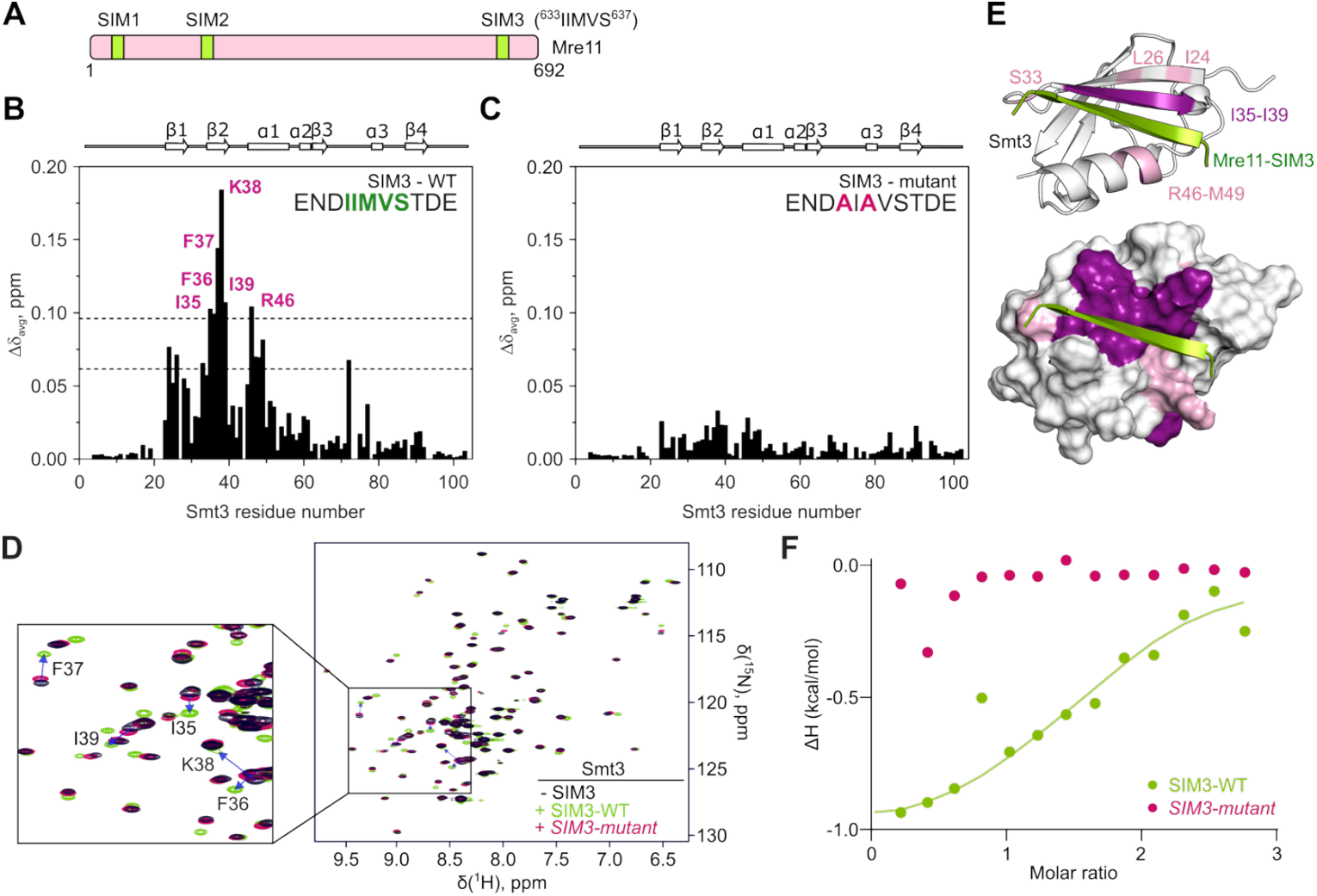
The Mre11 C-terminus contains a novel SUMO-interaction motif. **(A)** Schematic representation of SUMO-interacting motif (SIMs) in Mre11. **(B, C)** Average backbone amide chemical shift perturbations (Δδ_avg_) of Smt3 in the presence of 1.2 molar equivalents of (B) wild type or (C) mutant SIM peptide (sequences indicated). The secondary structure of Smt3 is shown above the plots. The horizontal lines in (B) correspond to the average Δδ_avg_ (avg) plus one or two standard deviations (stdev). **(D)** Overlaid [^1^H,^15^N] HSQC spectra of Smt3 in its free form (black) or with 1.2 molar equivalents of wild type (green) or mutant (pink) SIM3 peptides. Blue arrows indicate residues that experience highest chemical shift. **(E)** Chemical shift mapping of the wild-type SIM peptide binding. Smt3 is shown as (top) cartoon and (bottom) molecular surface coloured by the Δδ_avg_ values (pink: Δδ_avg_ > avg + stdev; violet: Δδ_avg_ > avg + 2*stdev). The bound SIM peptide is in green. The disordered regions at Smt3 N- and C-termini are omitted for clarity. **(F)** Normalized heat per peak ΔH (kcal/mol) as a function of molar ratio (peptide/protein concentration) measured by isothermal titration calorimetry upon titration of wild-type (green) or mutant (pink) SIM peptide on Smt3. The solid line shows the best fit to a single binding site model for the wild-type peptide.

To validate the Mre11-SIM3, we performed Nuclear Magnetic Resonance (NMR) spectroscopy on purified and isotopically labelled U-[^13^C, ^15^N] yeast Smt3 in the presence of a synthetic Mre11-SIM3 peptide (**Figures 6B and S7E**). As a control, we mutated residues corresponding to Mre11 I633 and M635 to alanine, predicted to contribute to the interaction with Smt3 (**Figure S7B**). NMR spectroscopy analysis indicated that the wild-type Mre11-SIM3 peptide induced strong backbone amide chemical shift to Smt3 residues I35, F36, F37, K38, I39 and R46, in contrast to the mutant peptide (**Figures 6B-6D**). Residues I35-I39 form part of the second β-sheet of Smt3 and R46 is located on the first α-helix that are both predicted to bind the SIM3 peptide (**Figure 6E**). In addition, analysis of methyl binding shifts closely agrees with the backbone amide chemical shift perturbation and further validate the AlphaFold-predicted model between Mre11-SIM3 and Smt3 (**Figure S8**).

To evaluate the affinity of Smt3 for Mre11-SIM3, we performed isothermal titration calorimetry (ITC) binding experiments between the Mre11-SIM3 peptides and Smt3. ITC analysis confirmed that the wild-type SIM3 peptide binds Smt3 with a K_D_ of 20 ± 10 μM, typical for SUMO-SIM interactions^49,50^. In contrast, the mutant SIM3 peptide failed to interact with Smt3 (**Figures 6F and S9**).

### SIM3 contributes to Mre11 recruitment and meiotic DSB formation

To test the role of Mre11-SIM3 *in vivo*, the core SIM3 motif (IIMVS) was substituted with alanines. *mre11-SIM3* mutant cells showed a small reduction in meiotic Mre11 foci (**Figure 7A**), which was associated with delayed and reduced DSB formation (**Figure 7B and S10C**). Combining the *SIM3* mutation with truncation of the Mre11 C-terminal α-helix (*mre11-ΔC15+SIM3*) further reduced Mre11 foci and abolished DSB formation (**Figures 7A and 7B**), while protein level remained unchanged (**Figures S10A and S10B**). Consequently, the *mre11-SIM3* mutant had slightly reduced spore viability (86%), while *mre11-ΔC15+SIM3* spores were completely inviable (**Figures 7C)**. The *mre11-SIM3* mutant also showed a minor delay in meiotic progression; while progression was accelerated in the *mre11-ΔC15+SIM3* mutant compared to the wild-type, likely due to the absence of DNA breaks **(Figure S10D)**. We conclude that SIM3 promotes the recruitment of Mre11 to the meiotic DSB machinery.

**Figure 7:**
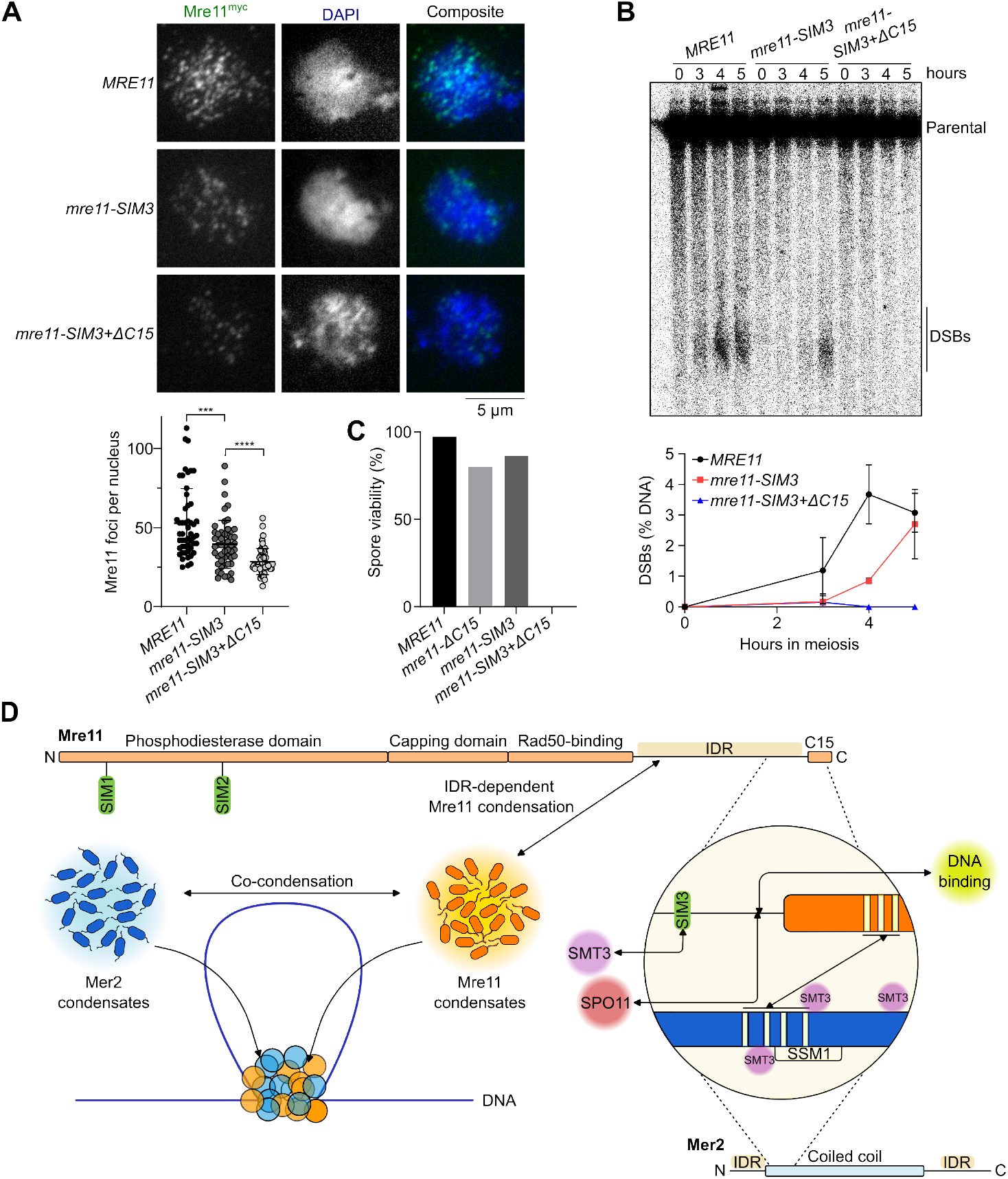
A SUMO-SIM interaction fosters the recruitment of Mre11 during meiosis. **(A)** Immunofluorescence on meiotic nuclear spreads of myc-tagged *MRE11-WT, mre11-SIM3*, and *mre11-SIM3+ΔC15* cells. Quantification of Mre11^myc^ foci show mean ± SD of n = 50 cells per strain. **(B)** Southern blot analysis of meiotic DSB formation at *GAT1* hotspot. Quantifications show mean ± range of n = 2 independent experiments. **(C)** Spore viabilities of wild-type and mutant *MRE11* strains (n ζ 88 spores). **(D)** Schematic model of the mechanisms that drive Mre11 recruitment during meiosis. Mre11 undergoes DNA-dependent condensation, driven by its IDR, and can mingle with Mer2 to form joint condensates. Mre11 is recruited to the DSB machinery via its C-terminus through a SUMO-SIM interaction and direct binding with Spo11 and Mer2.

To test the hypothesis that SIM3 contributes to the interaction between Mre11 and Mer2, we analyzed Mre11-Mer2 interaction using the yeast-two-hybrid assay. While truncation of the terminal 15 amino acids of Mre11 abolished the interaction with Mer2, mutation of the SIM3 motif had no discernable impact (**Figure S10E**). We note, however, that Mer2 may not be SUMOylated in this system, which employs mitotically cycling cells. Hence, the potential contribution of Mer2 SUMOylation to productive interaction with Mre11 during meiosis remains unclear.

## Discussion

### Budding yeast Mre11 has a C-terminal tail with meiosis-specific functions

The MRX complex plays key functions in the maintenance of genomic integrity throughout eukaryotes^51,52^. However, its role in promoting the formation of Spo11-dependent DSBs during meiosis has only been reported in budding yeast and *C. elegans*^15–17,53^. Indeed, MRX orthologs are not required for meiotic DSB formation in *A. thaliana*^54^, *S. pombe*^55^, and mice^56,57^. In *C. elegans*, MRE-11 and RAD-50 are necessary for DSB formation^53,58^, but the ortholog of Xrs2, NBS-1, is not^59^.

In *S. cerevisiae*, the binding of Mre11 to DSB hotspots depends on the presence of all other DSB proteins except Rad50^40^. This suggests that the MRX complex may be the final component recruited to the DSB machinery and raises the possibility that MRX recruitment may trigger Spo11’s catalytic activity, though the underlying mechanism remains unknown. It has been suggested that the requirement for the MRX complex prior to DSB formation serves to coordinate DSB formation with downstream repair, thereby minimizing genomic instability^40^. Supporting this idea, DSBs detected in wild-type cells are typically fully resected, indicating that processing occurs more rapidly than break accumulation. This observation is consistent with tight coordination between DSB formation and repair^60–63^.

It was previously shown that the C-terminus of Mre11 is dispensable for DSB repair in mitotically cycling cells, but essential for the formation of meiotic DSBs^29,40^. Here, we show that this essential meiotic function of the Mre11 C-terminus involves multiple mechanisms that collaborate to promote Mre11 recruitment to recombination sites (**Figure 7D**).

### The Mre11 C-terminus promotes meiotic DSB formation via multiple mechanisms

First, we show that the C-terminal α-helix of Mre11 directly binds the Mer2 SSM1 motif located on the N-terminal side of the tetrameric coiled coil. AlphaFold modeling and mutagenesis identified Mre11-LLK and Mer2-EQEK residues as being responsible for this interaction. Truncating the Mre11 terminal α-helix or mutating the Mer2-EQEK residues *in vivo* confirmed the functional importance of this interaction for Mre11 foci formation and DSB formation. These data are in line with previous yeast-two-hybrid and pulldown experiments that revealed an interaction between Mre11 and Mer2^41^. We note, however, that the Mer2 alleles previously reported to reduce Mre11 interaction involved amino acids predicted to point inside the Mer2 tetrameric coiled coil, which are therefore likely to affect Mre11 interaction indirectly by compromising the structural stability of Mer2 (**Figure S5C**).

Second, we identified and biochemically validated a novel SUMO-interacting motif located within the Mre11 IDR and showed that this motif also participates in Mre11 recruitment and DSB formation during meiosis. While the meiotic phenotypes of the *mre11-SIM3* mutant were relatively modest, we find that SIM3 collaborates with the C-terminal α-helix of Mre11 for productive recruitment to precursor DSB sites.

Third, a recent study demonstrated that the Mre11 C-terminus is also important for direct interaction with Spo11^31^. While the binding sites were not precisely identified, this interaction implicates Mre11 residues 663 to 676, just upstream of the terminal α-helix. It was noted that a mutant lacking the last 16 amino acids of Mre11 was defective in meiosis despite no effect on Spo11 binding, indicating that interaction with Spo11 is not sufficient for functional recruitment of Mre11. Our data explains this result by demonstrating that the Mre11 terminal α-helix binds to Mer2.

Finally, we showed that the C-terminal region of Mre11 is also required to assemble DNA-dependent condensates *in vitro*. DNA nucleates Mre11 condensates and acts as a scaffold that then recruits free soluble protein through homotypic self-association. We show that the C-terminal 49 residues of Mre11 are sufficient for self-association when Mer2 condensates are provided as a nucleation site, suggesting that this may constitute yet another meiosis-specific function of the Mre11 C-terminal tail. Nevertheless, in the absence of a mutant that specifically abolishes Mre11 self-association, the functional importance of this activity for meiotic DSB formation remains unclear.

In summary, the recruitment of Mre11 during meiosis involves at least three, perhaps four, independent functions of the Mre11 C-terminal tail.

### Assembly of the DSB machinery by hierarchical condensation

The *in vitro* condensation activity of Mre11 and MRX are reminiscent of that of Rec114-Mei4 and Mer2 that also form dynamic and reversible macromolecular assemblies in the presence of DNA^9^. Similar to RMM, Mre11 and MRX condensates likely involve electrostatic interactions between the negatively charged DNA backbone and positively charged residues present in the C-terminal region of Mre11. Condensates are further stabilized by the presence of positively charged ions such as magnesium that presumably inhibit intramolecular repulsion within the DNA substrate. In addition, sensitivity of Mre11 condensates to 1,6-hexanediol indicates that self-association via the low-complexity IDR depends on weak hydrophobic interactions, further supporting the liquid nature of these assemblies.

While both Mre11 and MRX condensates share similar properties, they don’t fully mirror each other in their behavior, suggesting that the presence of Rad50 and Xrs2 might confer additional stabilization to MRX condensates, consistent with a previous observation that Rad50 self-interaction drives Mre11-Rad50 oligomerization^34^.

The spontaneous assembly of Mer2, Rec114-Mei4 and Mre11/MRX condensates *in vitro* contrasts with the more stringent assembly of their respective foci *in vivo*. Indeed, Mer2 foci depend on the axis protein Hop1, Rec114 foci depend on Mer2, and Mre11 foci depend on Mer2 and likely most other DSB proteins^14,40,64–66^. We propose that the DSB machinery assembles via a hierarchical mechanism of successive condensate nucleation and growth events, where Hop1 nucleates the assembly of Mer2 condensates that nucleate Rec114-Mei4 condensates that recruit the Spo11 complex^9^. Mer2 condensates also recruit Mre11 and drive its self-assembly, dependent on stabilizing interactions with Spo11 and other SUMOylated targets (**Figure 7D**).

### The importance of a SUMO-SIM interaction in Mre11 recruitment

If the recruitment of Mre11 constitutes the final step in the assembly of the DSB machinery prior to triggering Spo11-dependent DNA cleavage, SUMOylation of the DSB machinery could serve to mark licensed pre-DSB complexes and/or allow for reversible interactions.

What are the SUMOylated targets bound by Mre11 prior to DSB formation? SUMOylation was previously shown to regulate the key events of meiotic prophase I, including DSB formation, and thousands of SUMOylation sites were mapped on meiotic proteins^45^. Amongst those, thirteen sites were identified within Mer2, including several close to the SSM1 motif. Given the physical proximity of the Mre11^LLK^ and Mer2^EQEK^ interaction regions and Mre11-SIM3, it is tempting to speculate that SIM3 might be interacting with SUMOylated Mer2 during meiosis, which would presumably serve as an anchor to stabilize the interaction with Mer2. However, the recruitment of Mre11 through the SIM3 motif may involve other SUMOylated DSB proteins, including Spp1, Rec114, Hop1, Red1, and cohesin^45^.

## Materials and Methods

### Preparation of expression vectors

Oligonucleotides (oligos) used in this study were purchased from Sigma-Aldrich. The sequences of the oligos used are listed in **Table S1**. Plasmids generated in this study were verified by sequencing and are listed in **Table S2**. Peptides used were ordered from GenScript or synthesized in the Ballet laboratory and are listed in **Table S3**.

The expression vector for Mre11^10xHis^ was produced by PCR amplification of the *MRE11* gene from yeast genomic DNA (SK1 strain) using primers cb1351 and cb1352 and Gibson assembly into a BamHI and EcoRI digestion fragment of pFastBac1 to yield pCCB865. The sequence coding for eGFP was cloned into the BamHI site of pCCB865 to produce the expression vector for ^eGFP^Mre11, pCCB942. The expression vector for Rad50 was produced by PCR amplification of the *RAD50* gene from SK1 genomic DNA using primers cb1353 and cb1354 and Gibson assembly into a BamHI and EcoRI digestion fragment of pFastBac1 to yield pCCB866. The expression vector for Xrs2^2xFlag^ was produced by PCR amplification of the *XRS2* gene from SK1 genomic DNA using primers cb1355 and cb1356 and Gibson assembly into a BamHI and EcoRI digestion fragment of pFastBac1 to yield pCCB867.

Expression vectors for ^eGFP^Mre11^10xHis^-*Δ10-270* (pry7), ^eGFP^Mre1^10xHis^-*Δ290-472* (pry5), ^eGFP^Mre1^10xHis^-Δ*IDR* (pry6), and ^eGFP^Mre1^10xHis^-Δ*C49* (pCCB943) were amplified by inverse PCR from pCCB942 using primers pp26 and pp27, pp28 and pp29, pp24 and pp25, and cb1435 and cb1458, respectively. The amplified product was gel extracted, phosphorylated, and ligated to generate the truncations. The expression vector for ^eGFP^Mre11-IDR (pry41) was generated by Gibson assembly using a backbone amplified from pCCB942 with primers pp55 and pp61 and the Mre11-IDR sequence amplified from pCCB942 using primers pp59 and pp60.

The vector for expression of ^MBP^Mre11-C49 and ^HisSUMO^Mer2 (pCCB1040) was based on a pET-Duet1 vector with the sequence coding for ^MBP^Mre11-C49 cloned within the first position (SacI site) and the sequence coding for ^HisSUMO^Mer2 cloned at the second position (XhoI site). The EQEK mutations and LLK mutations were introduced by PCR amplification of pCCB1040 with primers cb1561 and cb1562, and pp84 and pp85, followed by phosphorylation and self-ligation to yield plasmids pry59 and pry61, respectively.

The expression vector for ^mScarlet^Mre11-C49 (pry109) was generated by performing a three-fragment Gibson assembly of a pET28a backbone containing MBP followed by a TEV cleavage site amplified from pCCB785 using primers cb1486 and pp120, Mre11-C49 amplified from pCCB1040 using primers pp92 and pp136, and mScarlet amplified from pCCB785 using primers pp148 and pp149. Expression vectors for ^Alexa594^Mer2 (pCCB750) and ^eGFP^Mer2 (pCCB777) were previously described^9^.

The pET28b expression vector for Smt3 (pCCB998) was a kind gift from Chris Lima^67^. Plasmids for yeast 2-hybrid, pWL1592 (pGBDU-C1-Mer2), pWL1596 (pGAD-C1-Mre11), and pWL1565 (pGAD-C1) were generously provided by John Weir. Mutations in the coding region of Mre11 were introduced in pWL1596 via PCR mutagenesis using the primers RB70 and RB267 to generate pGAD-C1-Mre11-*SIM3* (pNH1371) and RB268 and RB269 to generate pGAD-C1-Mre11-*ΔC15* (pNH1372).

### Expression and purification of recombinant proteins

Recombinant baculoviruses were produced by Bac-to-Bac Baculovirus Expression System (Invitrogen) following the manufacturer’s instructions. For every induction, 1 L culture containing 2 × 10^6^ *Spodoptera frugiperda* (Sf9) cells/ml were infected with a Multiplicity of Infection (MOI) of 2.5 for each of the viruses. Viruses generated from pCCB865, pCCB866, and pCCB867 were used for expression of Mre11^10xHis^-Rad50-Xrs2^2xFLAG^ (MRX) and pCCB942, pCCB866, and pCCB867 were used for the expression of ^eGFP^Mre11^10xHis^-Rad50-Xrs2^2xFLAG^ (^eGFP^MRX). Full-length Mre11^10xHis^ and ^eGFP^Mre11^10xHis^ were expressed using viruses generated from pCCB856 and pCCB942, respectively. Mre11 truncations ^eGFP^Mre11^10xHis^-*Δ10-270*, ^eGFP^Mre11^10xHis^-*Δ290-472*, ^eGFP^Mre11^10xHis^-*ΔIDR*, and ^eGFP^Mre11^10xHis^-*ΔC49* were expressed using viruses generated from pry7, pry5, pry6, and pCCB943, respectively.

Prior to harvest, Sf9 cells were allowed to infect for 62 hours at 27 ºC at 80 rpm, following which cells were pelleted at 500 rcf, washed once with 1x PBS, snap frozen in liquid nitrogen, and stored at -80ºC, or used for purification. All subsequent purification steps were carried out at 0 - 4 ºC.

The following protocol was used for the purification of His-tagged ^eGFP^Mre11 and ^eGFP^Mre11 truncations: Frozen pellets were resuspended in lysis buffer (25 mM HEPES, pH 7.5, 20 mM imidazole, 0.1 mM DTT, Roche Complete Tablet (11836170001), and 0.3 mM PMSF) and made up to a total volume of 35 ml. The samples were transferred to a beaker and osmotic lysis was performed by slowly adding 5 ml of 5 M NaCl (final 500 mM) and 10 ml of 50% (vol/vol) glycerol (final 10%) while gradually mixing with a stir bar for 30-40 mins. Lysed cells were centrifuged at 30,000 rpm for 30 mins and soluble fraction was used for affinity chromatography. 1 ml Ni-NTA resin (Thermo Scientific, 88223) was pre-equilibrated with wash buffer (25 mM HEPES, pH 7.5, 500 mM NaCl, 10% glycerol, 20 mM imidazole, 0.1 mM DTT, 0.3 mM PMSF) and batch incubated with the soluble fraction for 1 h. The resin was washed extensively in wash buffer and eluted with wash buffer containing 500 mM imidazole. Peak fractions were pooled and loaded onto a Superdex 200 Increase 10/300 GL column pre-equilibrated with SEC buffer (25 mM HEPES, 10% glycerol, 1 mM DTT, 2 mM EDTA, 300 mM NaCl). Following size-exclusion chromatography, fractions containing protein were concentrated using a 30 kDa MWCO Amicon ultra centrifugal filters (Millipore), aliquoted, snap froze in liquid nitrogen, and stored at -80°C.

For fluorescence labelling of Mre11, the Ni-NTA eluate was dialyzed several times to remove traces of imidazole. Labeling reaction was performed using Alexa Fluor 488 Protein Labeling Kit (Invitrogen, A10235) that has a succinimidyl ester moiety that reacts with primary amines. After 1 hour conjugation at room temperature, unbound fluorophore was removed by size-exclusion chromatography as described above.

Purification of recombinant His- and Flag-tagged ^eGFP^MRX complexes and truncations were performed essentially as described^68^. Briefly, following osmotic lysis, soluble extract was used for sequential affinity chromatography with Ni-NTA resin (Thermo Scientific, 88223) and anti-FLAG M2 affinity gel (Sigma, A2220). Peak eluted fractions were pooled, aliquoted, and snap frozen.

For expression of recombinant ^mScarlet^Mre11-C49 in *E. coli*, pry109 was transformed in BL21 cells and plated on LB plates containing kanamycin. Cells were then cultured in LB media at 37°C to an optical density (OD_600_) of 0.6. Expression was carried out for 20 hours at 16°C with 0.3 mM isopropyl β-D-1-thiogalactopyranoside (IPTG) following which cells were pelleted at 3000 rcf, washed once with 1x PBS, snap frozen in liquid nitrogen, and stored at -80ºC, or used for purification. Cell pellet was resuspended in lysis buffer containing 25 mM HEPES, 500 mM NaCl, 0.1 mM DTT, 0.01% NP40, 10% glycerol, 1 mM PMSF, Protease Inhibitor Cocktail (PIC, 1:100, Sigma), and 2 mM EDTA. Cells were lysed by sonication (15 W, 5 mins, 5 sec pulse) and centrifuged at 20,000 rpm for 20 mins. Soluble extract was incubated for 1 hour with 1.5 ml amylose resin (E8021L, NEB), pre-equilibrated with wash buffer (25 mM HEPES, 500 mM NaCl, 0.1 mM DTT, 0.01% NP40, 10% glycerol, 0.5 mM PMSF, Protease Inhibitor Cocktail (PIC, 1:200, Sigma), 2 mM EDTA). The column was washed extensively with wash buffer and then elution was performed in wash buffer containing 10 mM maltose. Peak fractions were pooled, the MBP-tag was cleaved with TEV protease overnight without rotation and then loaded on a Superdex 75 Increase 10/300 GL column pre-equilibrated with SEC buffer (25 mM HEPES, 300 mM NaCl, 10% glycerol, 2 mM EDTA). Fractions containing protein were concentrated in 10 kDa MWCO Amicon ultra centrifugal filters (Millipore), aliquoted, snap froze in liquid nitrogen, and stored at -80°C.

Expression and purification of ^eGFP^Mer2 and ^Alexa594^Mer2 were performed as previously described^9^.

For expression of recombinant Smt3^6xHis^ in *E. coli*, pCCB998 was transformed in BL21 (DE3)pLysS cells and plated on LB plates containing kanamycin. Cells were then cultured in LB media at 37°C to an OD_600_ of 0.6. Expression was carried out for 3 hours at 37°C with 1 mM IPTG. Cells were centrifuged at 18°C at 4000 g for 15 mins and were directly resuspended in lysis buffer (20 mM NaPi, pH 6.5, 30 mM imidazole, 350 mM NaCl, 0.1 mM DTT, 1 mM PMSF). Cells were lysed by sonication (10 W, 4 mins, 4 sec pulse) and centrifuged at 20,000 rpm for 20 mins. Soluble fraction was incubated for 1 hour with 1.5 ml Ni-NTA resin, pre-equilibrated with wash buffer (20 mM NaPi, pH 6.5, 30 mM imidazole, 350 mM NaCl, 0.1 mM DTT, 0.1 mM PMSF). The column was washed extensively with wash buffer and then eluted in wash buffer containing 500 mM imidazole.

For the production of the doubly labeled U-[^13^C, ^15^N] Smt3^6xHis^ protein, the IPTG induction was carried out in minimal medium containing M9 salts (6.8 g/L Na_2_HPO_4_, 3 g/L KH_2_PO_4_, and 1 g/L NaCl), 2 mM MgSO_4_, 0.2 mM CaCl_2_, trace elements (60 mg/L FeSO_4_·7H_2_O, 12 mg/L MnCl_2_·4H_2_O, 8 mg/L CoCl_2_·6H_2_O, 7 mg/L ZnSO_4_·7H_2_O, 3 mg/L CuCl_2_·2H_2_O, 0.2 mg/L H_3_BO_3_, and 50 mg/L EDTA), BME vitamin mix (Sigma), and 1 g/L ^15^NH_4_Cl and 2 g/L [^13^C_6_]glucose (CortecNet) as the sole nitrogen and carbon sources, respectively. The purification protocol remained the same as described above.

### In vitro condensation assays

Proteins were diluted to a 10× stock of their appropriate working concentrations in their respective storage buffers. Reactions were performed in a buffer containing 25 mM HEPES-HCl (pH 7.5), 2 mM DTT, 1 mg/ml BSA, 5% glycerol, 5 mM MgCl_2_, 5% PEG-3350, and NaCl. Considering the salt contributed by the protein dilution buffer, final concentration of NaCl in a reaction was adjusted to 120 mM. Unless specified otherwise, all reactions contained 400 nM ^Alexa488^Mre11 or 100 nM ^eGFP^MRX. A typical 20 μL binding reaction contained 2 μL protein of 10× stock of indicated concentration, 10 μL of 2× reaction buffer, and 150 ng of supercoiled pUC19 (5.7 nM). Typical reactions were assembled at 30°C for 30 mins with gentle mixing every 5 mins, unless mentioned otherwise. 5 μL was dropped onto a microscope slide and covered with a coverslip. All images were captured on a Zeiss Axio Observer with a 100×/1.4 NA oil immersion objective except for images provided in Figure 4G which were captured on Leica Stellaris DMI 8 confocal microscope with a 63x/1.2 NA water immersion objective. Images were analyzed with ImageJ using a custom-made script^9^. In brief, 129.24 × 129.24-μm (2048 × 2048-pixel) images were thresholded to mean intensity of the background plus three times the standard deviation of the background. Masked foci were counted and the intensity inside the focus mask was integrated. Data points represent averages of at least 8-10 images per sample. Data were analyzed using Graphpad Prism 10.4.0.

### Gel shift assays

Proteins were diluted to their appropriate working concentrations in their respective storage buffers. A typical 20 μL binding reaction was performed in a reaction buffer containing 25 mM HEPES-HCl (pH 7.5), 2 mM DTT, 1 mg/ml BSA, 10% glycerol, 5 mM EDTA, and NaCl adjusted to a final concentration of 100 mM, 1 nM pUC19 plasmid substrate, and the indicated concentration of protein. Reactions were assembled at 30°C for 30 mins and resolved in a 1% agarose (SeaKem LE Agarose, Lonza) at 60 V for 120 mins at 4°C. Gels were stained with SYBR Gold Nucleic Acid Gel Stain (S11497, Invitrogen) for 40 mins and visualized with Amersham Typhoon biomolecular imager (Cytiva).

### Yeast targeting vectors and strain construction

Yeast strains are generated from *Saccharomyces cerevisiae* SK1 background and are listed in **Table 4**.

To produce a yeast targeting vector, *MRE11-myc8::URA3* was amplified using cb1424 and cb1425 from the genomic DNA of CBY375 (SKY1361) and cloned into TOPO vector by TOPO blunt cloning, generating pry2. Plasmids to produce *mre11-ΔIDR* (pry30), *mre11-ΔC49* (pry24), and *mre11-ΔC15* (pry42) mutants were generated by inverse PCR followed by self-ligation of pry2 using primers pp24 and pp25, cb1435 and pp46, and pp72 and pp73, respectively. Plasmids for the *mre11-SIM* (pry56) and *mre11-SIM+ΔC15* (pry57) were generated similarly using primers pp3 and pp4 and templates pry2 and pry42, respectively. Genomic integration of wild-type and truncated *MRE11-myc8::URA3* cassettes was performed by SpeI and NotI digestion of the corresponding plasmids and insertion in the endogenous *MRE11* locus of strain CBY006 by ‘LiAc’-based transformation.

Plasmids to produce *MER2* mutant strains were based on pMH002, which contains a *MER2::HphMX* cassette cloned into a TOPO-based vector, as described^43^. To generate *mer2-EQEK*, pMH002 was amplified by inverse PCR followed by ligation using primers cb1561 and cb1562 to yield pCCB1046. An internal V5 (iV5) tag was introduced between Mer2 residues 248 and 249 (Mer2^iV5^) using primers dam005 and dam006 on plasmid backbone pMH002 by inverse PCR followed by ligation to generate pDAM003. *mer2*^*iV5*^*-EQEK* was constructed by inverse PCR followed by ligation of pDAM003 using primers cb1451 and cb1562. Genomic integration of *mer2-EQEK::HphMX* and *mer2*^*iV5*^*-EQEK::HphMX* cassettes was performed by SpeI, NotI, and XmaI digestion of the respective plasmids and insertion into the endogenous *MER2* locus of CBY006 by ‘LiAc’-based transformation. The *MER2*^*iV5*^::*HphMX* allele was constructed similarly following BamHI and SphI digestion of pDAM003.

All strains were genotyped by PCR and sequencing and opposite mating type was generated by crossing with CBY007. All other yeast strains were generated by crossing with appropriate genotypes listed in **Table S4**.

### Spore viability assay

For spore viability assay, a small patch of diploid strain was incubated in sporulation media (2% potassium acetate) at 30°C, 250 rpm for two days. After two days, 1 mL of sporulating culture was centrifuged and all but 200 μL of supernatant was removed. To digest yeast cells, 2 μL of concentrated sporulation culture was mixed with 100 μL 1 M sorbitol and 1 μL 10 mg/ml zymolyase and incubated at 30°C for 21 mins. 20 μL of digested cells were dropped on a YPD plate, left to dry for about 10 mins, and were micromanipulated using a tetrad dissector (MSM400, Singer Instruments). At least 20 tetrads were dissected for each assay and spore viability by assessed by calculating the number of viable spores after 2 days of incubation at 30°C.

### Yeast culture and meiotic synchronization

Following standard protocols, strains were patched on YPG plates, mated and streaked on YPD plates, and selected diploid colonies grown in liquid YPD at 30°C, 250 rpm, overnight. For meiotic synchronization, diploid cultures grown overnight in YPD were transferred to YPA (1% yeast extract, 2% peptone, 1% potassium acetate) at OD_600_ 0.2 and grown for 12-14 hours at 30°C, 250 rpm. Once the cultures reached OD_600_ between 1.2-1.6, cells were washed once with prewarmed sterile water and immediately transferred to sporulation medium supplemented with amino acids (320 μL amino acid complementation media for 100 mL of sporulation media (SPM)) and were kept shaking at 30°C, 250 rpm during the entire meiotic time-course. For MMS and CPT (Sigma) sensitivity assays, serial dilutions of overnight cultures were spotted on freshly prepared YPD-MMS or YPD-CPT plates containing indicated percentage of MMS or CPT, respectively. Plates were grown for two days at 30°C. For immunofluorescence of vegetatively growing cells, overnight cultures were refreshed by diluting to OD_600_ 0.2 and grown for 3-4 hours to reach OD_600_ 1.2-1.4 before subjecting to MMS or MMS followed by 5% 1,6-hexanediol treatment. For 5% 1,6-hexanediol treatment, cells were first converted to spheroplasts and then treated with 5% 1,6-hexanediol for 4-5 minutes. Spheroplasts were then immediately washed, lysed, and fixed using protocol described below.

### Spreading and immunofluorescence of yeast nuclei spreads

Meiotic cultures were harvested 4 hours after transfer to SPM, washed with Recruitment of Mre11 during meiosis sterile cold water, and resuspended in 1 M sorbitol, 1× PBS (pH 7), 10 mM DTT, 0.5 mg/ml zymolyase 20T, and incubated for 30 mins at 30°C with gentle shaking. Spheroplasts were collected by centrifuging for 1 min at 1500 rpm and were washed gently with ice-cold 0.1 M MES-1 M sorbitol. Spheroplasts were then centrifuged, lysed by adding ice-cold 0.1 M MES and 4% paraformaldehyde followed by vigorous finger-vortexing and immediately fixing on microscopy slides for 1 hour at room temperature. Slides were washed three times with 1 ml 0.4% PhotoFlo 200 solution (Kodak), air dried and stored at -20°C or directly used for processing.

Slides were blocked with 90% FBS, 1× PBS for 1 hour at room temperature in a humid chamber and then incubated with primary antibody (mouse mAb anti-myc, 1:200 (2276S, Cell Signaling Technology), rabbit anti-phospho H2A-S129, 1:200 (ab15083, abcam)) diluted in 3% BSA, 1× PBS in a humid chamber for 2 hours at 37°C or overnight at 4°C. Slides were washed three times with 1× PBS in a Coplin jar, were incubated with secondary antibody (goat anti-mouse IgG Alexa Fluor™ Plus 488, 1:200 (A32723, Invitrogen), donkey anti-rabbit IgG Alexa Fluor™ 546, 1:200 (A10040, Invitrogen) diluted in 3% BSA, 1× PBS in a humid chamber at 37°C for 1 hour. Slides were washed in the dark three times for 5 mins with 1× PBS, mounted with Vectashield containing DAPI (Vector Labs). Images were captured on a Zeiss Axio Observer with a 100×/1.4 NA oil immersion objective and were analyzed in ImageJ.

### Western blotting of yeast meiotic extracts

Meiotic cultures at desired timepoints were harvested, washed in ice-cold water, and lysed in 20% trichloroacetic acid (TCA) by agitation in a bead beater (insert company) using 0.5 mm zirconia/silica beads (insert company). Precipitated proteins were solubilized in Laemmli sample buffer and appropriate amounts of protein were separated by SDS-PAGE and analyzed by Western blotting. Western blotting was performed using standard protocol. Primary antibody used was mouse monoclonal anti-myc at 1:1000 dilution (2276S, Cell Signaling Technology), mouse monoclonal anti-V5 at 1:500 (R96025, Invitrogen), mouse monoclonal anti-PGK1 at 1:5000 (ab113687, Abcam) and secondary antibody used was goat anti-mouse IgG-HRP conjugated at 1:10,000 dilution (AP308P, Chemicon). Western blots were revealed using SuperSignal West Femto Maximum Sensitivity Substrate (Thermo Scientific) in Amersham Imager 600 (Cytiva).

### Southern Blotting

Meiotic DSB analysis by Southern blotting was performed as previously described^69^. In brief, synchronized cultures undergoing meiosis were collected at the indicated time points. After DNA isolation, 1 μg of genomic DNA was digested by PstI and separated on a 1% TBE-agarose gel. DNA was transferred to Amersham™ Hybond™-N+ nylon membranes (Cytiva) by vacuum transfer, hybridized with *GAT1* probe (amplified with primers: 5′-CGCGCTTCACATAATGCTTCTGG, 5’-TTCAGATTCAACCAATCCAGGCTC) and developed by autoradiography.

### Pull-down assay

Wild-type and mutant MBP-tagged Mre11-C49 and HisSUMO-tagged Mer2 were co-expressed in 50 mL of *E. coli* BL21 cultures and purified by affinity chromatography on Ni-NTA resin following a procedure similar to that described above. Briefly, cells were lysed by sonication and centrifuged at maximum speed for 30 mins at 4°C on a table-top centrifuge. A small fraction of the supernatant was collected as ‘Input’ and the remainder was incubated with 120 μL Ni-NTA resin, pre-equilibrated with wash buffer (25 mM HEPES pH 7.5, 20 mM imidazole, 0.1 mM DTT, 0.1 mM PMSF, 10% glycerol), for 1 hour on a rotating wheel at 4°C. The resin was washed twice with 2 ml in batch and five times with 2 ml on column before eluting with 250 μL of 500 mM imidazole in wash buffer. Input and elution fractions were separated by SDS-PAGE followed by immunoblotting with primary murine anti-MBP monoclonal antibody at 1:10,000 dilution (E8032S, NEB) and secondary goat anti-mouse IgG-HRP conjugated at 1:10,000 dilution (AP308P, Chemicon). Western blots were revealed using SuperSignal West Femto Maximum Sensitivity Substrate (Thermo Scientific) in Amersham Imager 600 (Cytiva).

### Nuclear magnetic resonance spectroscopy

The samples contained 0.3-0.6 mM of U-[^13^C,^15^N] Smt3 in 20 mM sodium phosphate, 20 mM NaCl (pH 6.5), 0.02% NaN_3_ and 10% D_2_O for the lock. All NMR spectra were acquired at 298 K on a Bruker Avance III HD 800 MHz spectrometer, equipped with a TCI cryoprobe. The NMR data were processed in TopSpin 3.6 (Bruker) or NMRPipe^70^ and analyzed in CCPNMR^71^. Assignments of Smt3 backbone amide resonances were taken from literature^72^ and verified by 3D HNCACB, HN(CO)CACB, and ^15^N-edited NOESY-HSQC spectra. Further assignments of methyl resonances were obtained from 3D HBHA(CO)NH and (H)CCH TOCSY experiments performed on the wild-type SIM-bound Smt3 sample, which exhibited superior spectral quality compared to that of the free protein. The assigned ^1^H, ^13^C and ^15^N chemical shifts of the free and bound Smt3 have been deposited in the Biological Magnetic Resonance Bank (http://www.bmrb.wisc.edu/) under the accession number 53209.

The NMR binding experiments were performed by an incremental addition of wild-type or mutant SIM peptides (1.2 mM stocks in the working buffer) to 0.3 mM samples of U-[^13^C,^15^N] Smt3, with the spectral changes monitored in [^1^H,^15^N] HSQC spectra acquired at each increment. The average chemical shift perturbations (Δδ_avg_) were calculated as Δδ_avg_ = (Δδ_X_^2^/50 + Δδ_H_^2^/2)^0.5^, where Δδ_X_ and Δδ_H_ are the chemical shift changes of the backbone amide nitrogens or methyl carbons (Δδ_X_) and protons (Δδ_H_) of Smt3 residues upon addition of 1.2 molar equivalents of SIM peptides, and n = 50 or 9 for the backbone amide and methyl groups, respectively.

### Peptide synthesis

The peptides were synthesized by standard Fmoc-based solid phase peptide synthesis. Preloaded Fmoc-Glu-Wang resin (0.55 mmol/g) was swollen in *N,N*-dimethylformamide (DMF) and the Fmoc-deprotection steps consisted of shaking the resin in two consecutive steps of 5 minutes and 15 minutes, in 20% 4-methylpiperidine in DMF containing 0.1 M 1-hydroxybenzotriazole (HOBt). The coupling steps were performed with conventional Fmoc-protected amino acids (3 equiv.) (except for N-α-Fmoc-(O-3-methyl-pent-3-yl)aspartic acid), hexafluorophosphate benzotriazole tetramethyl uronium (HBTU, 3 equiv.) and *N,N*-diisopropylethylamine (6 equiv.). After synthesis of the full sequence on resin, the peptide was cleaved with trifluoroacetic acid (TFA)/triisopropylsilane (TIS)/H_2_O (90/5/5, *v/v*). The products were purified by preparative reverse phase HPLC using an acetonitrile/water eluent mixture containing 0.1% TFA.

### Isothermal titration calorimetry

The measurements were performed in 20 mM NaPi 100 mM NaCl pH 7.5 on Microcal ITC200 calorimeter at 25°C. The syringe contained 2 mM of peptide, while the cell held 150 μM or 300 μM of protein. Given the poor peptide solubility in aqueous buffers, the peptide solutions necessitated addition of 10% DMSO. For the ITC titrations, equal amounts of DMSO (10%) were included in both compartments to minimize the heat of dilution. Each titration consisted of a first injection of 0.4 μL, followed by 12-13 injections of 2 μL peptide into the cell, separated by intervals of 120 seconds. The first injection was discarded during the analysis of the data. The wild-type peptide titration on Smt3 was performed in duplicate. The Microcal LLC ITC200 Origin software was used to fit the data to a single binding site model.

### Yeast two-hybrid

Bait and prey plasmids were co-transformed into yWL365 using the standard ‘LiAc’-based transformation and plated on SC-Leu-Ura selective media. At least four independent transformants were tested for Mer2-Mre11 interaction by spotting a dilution series on SC-Leu-Ura-His + 25 mM 3-AT (3-amino-1,2,4-triazole) and growing for 4-5 days at 30°C.

### Statistical analysis and data visualization

All statistical analysis and graphing were performed using Graphpad Prism version 10.4.0. Student’s t-test was used for determination of statistical significance and *P*-value calculation (p ≥ 0.05, ns, not significant; \*\**p* < 0.01; \*\*\**p* < 0.001; \*\*\*\**p* < 0.0001).

## End Matter

### Author Contributions and Notes

P.P. designed, executed, and analyzed all experiments except as noted; M. S. performed Southern blot experiments; W.E.Y.M. synthesized peptides under the supervision of S.B., performed ITC experiments and analyzed NMR data; A.N.V. acquired and analyzed NMR data; R.B. performed yeast 2-hybrid experiments under the supervision of N.H.; C.C.B. supervised the research and secured funding. P.P. and C.C.B wrote the paper with input from all authors. The authors declare no conflict of interest.

## Acknowledgments

We thank David Alvarez Melo for generating Mer2-iV5 tagged strains, John Weir for plasmids and strains, and CCB laboratory members for discussion. We also thank Biological Imaging facility (IMABIOL) at UCLouvain and Marie-Christine Eloy for providing training in the use of the epifluorescence microscope. This work was supported by the European Research Council under the European Union’s Horizon 2020 research and innovation program (ERC grant agreement 802525 to CCB), and the Fonds National de la Recherche Scientifique (PDR grant T.0031.22 to CCB). PP is funded by FNRS Aspirant fellowships (project 1.A908.22). CCB is a FNRS Research Associate. WEYM and SB acknowledge the Research Council of VUB for support through the Strategic Research Program SRP95 and the infrastructure grant OZR3939. NIH NIGMS grant R01GM074223 supported NH, who is also an Investigator of the Howard Hughes Medical Institute.

## Supplementary Information

**Figure S1:**
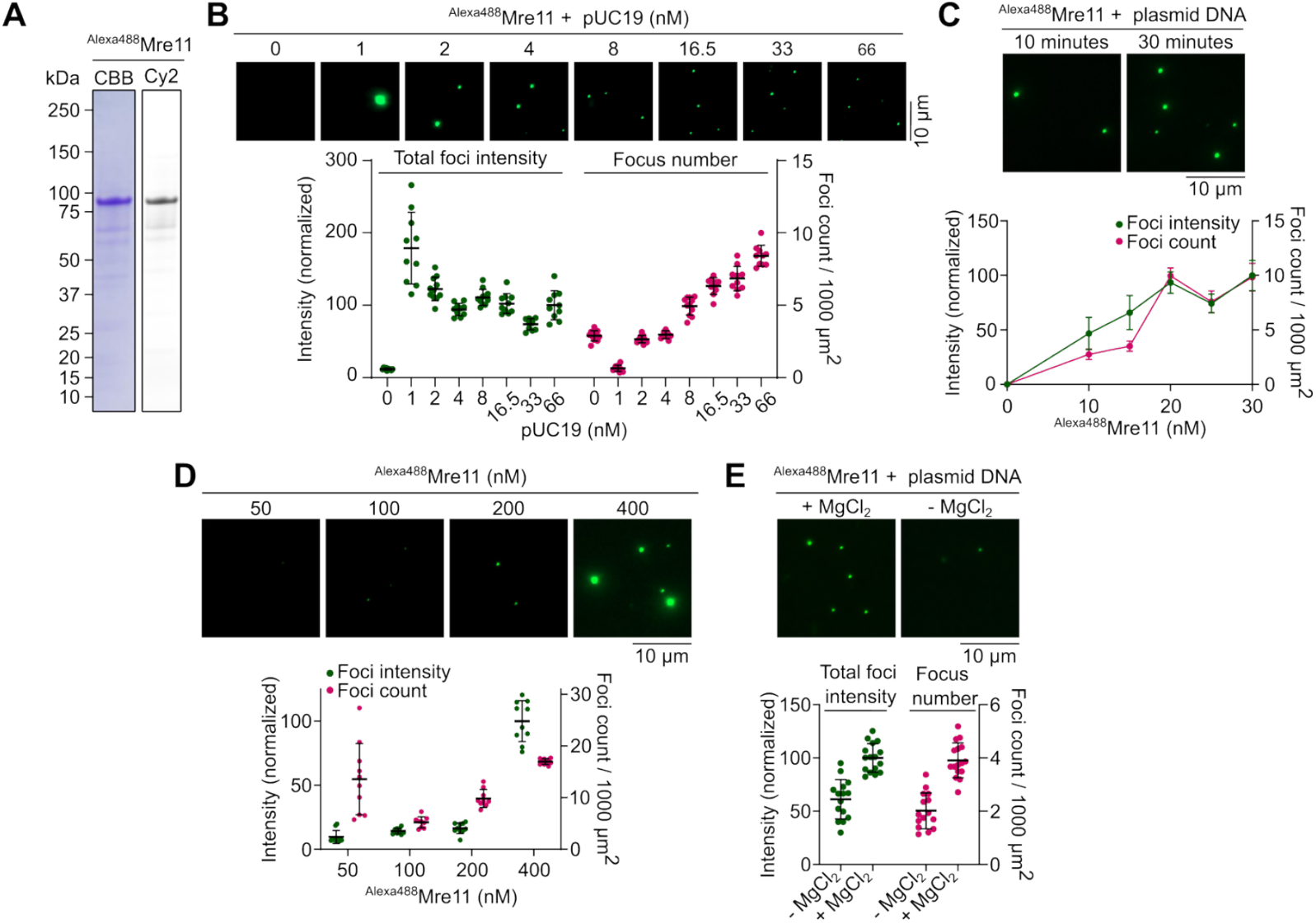
Properties of Mre11 nucleoprotein condensates. **(A)** SDS-PAGE of Alexa488-labelled Mre11 stained with Coomassie Brilliant Blue (CBB) (left) and visualized with a Cy2 filter (right). **(B)** Effect of plasmid DNA (pUC19) concentration on ^Alexa488^Mre11 condensate, visualized by epifluorescence microscopy. Quantification shows total fluorescence intensity (green) in a field of view normalized to the highest DNA concentration, and total number of foci per 1000 μm^2^ (magenta). Error bars represent mean ± SD from 10-11 fields of view. **(C)** Time-dependent change in Mre11 condensate assembly. Reactions contained 400 nM ^Alexa488^Mre11, 5.7 nM plasmid DNA and 5% PEG. Samples were collected at the indicated time points, immediately placed on a glass slide, covered with coverslip, and imaged. Foci intensities are normalized to the mean of the sample drawn at 30 minutes post-incubation. Error bars represent mean ± SD from 10-15 fields of view. **(D)** Effect of Mre11 concentration on condensation in presence of plasmid substrate and 5% PEG. Foci intensities are normalized to the mean of the sample with 400 nM Mre11. Error bars represent mean ± SD from 10 fields of view. **(E)** Effect of presence of divalent cation (5 mM MgCl_2_) on Mre11 condensates assembled in presence of 5.7 nM pUC19. Foci intensities are normalized to reaction in the presence of magnesium. For experiment performed without magnesium (E), 5 mM EDTA was included in the reaction.

**Figure S2:**
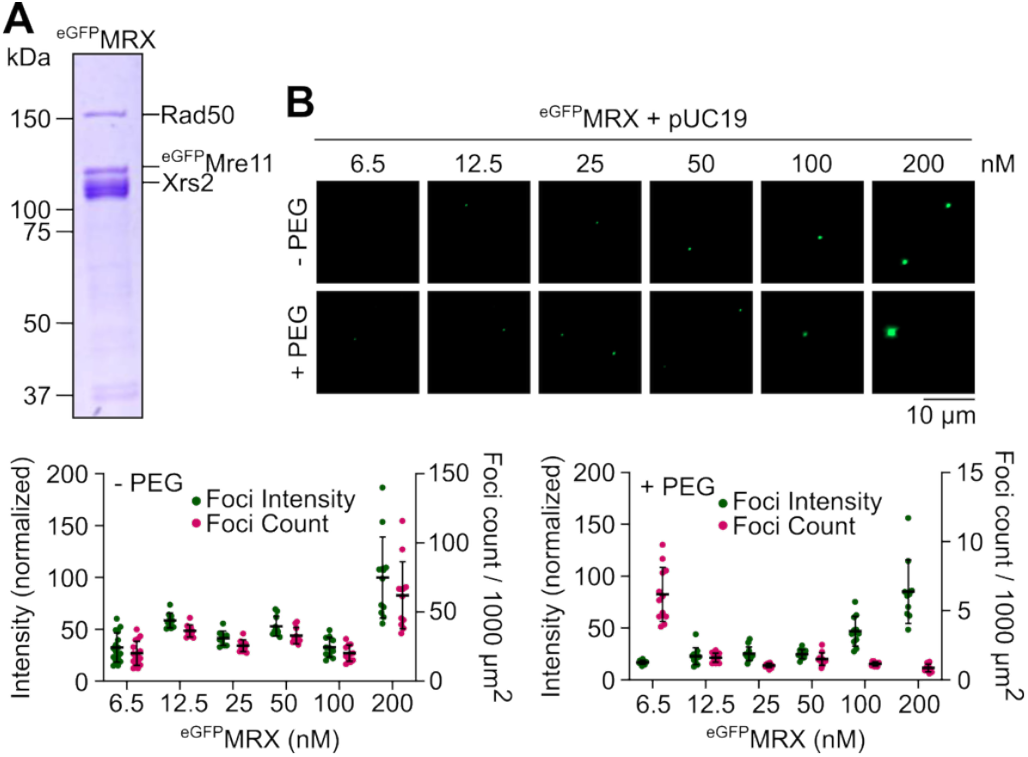
Properties of MRX nucleoprotein condensates. **(A)** SDS-PAGE of eGFP-tagged Mre11-Rad50-Xrs2 (MRX) complex stained with Coomassie Brilliant Blue. **(B)** Effect of MRX concentration on nucleoprotein condensation in the presence or absence of 5% PEG. Foci intensities are normalized to the mean of the sample with 400 nM Mre11.

**Figure S3:**
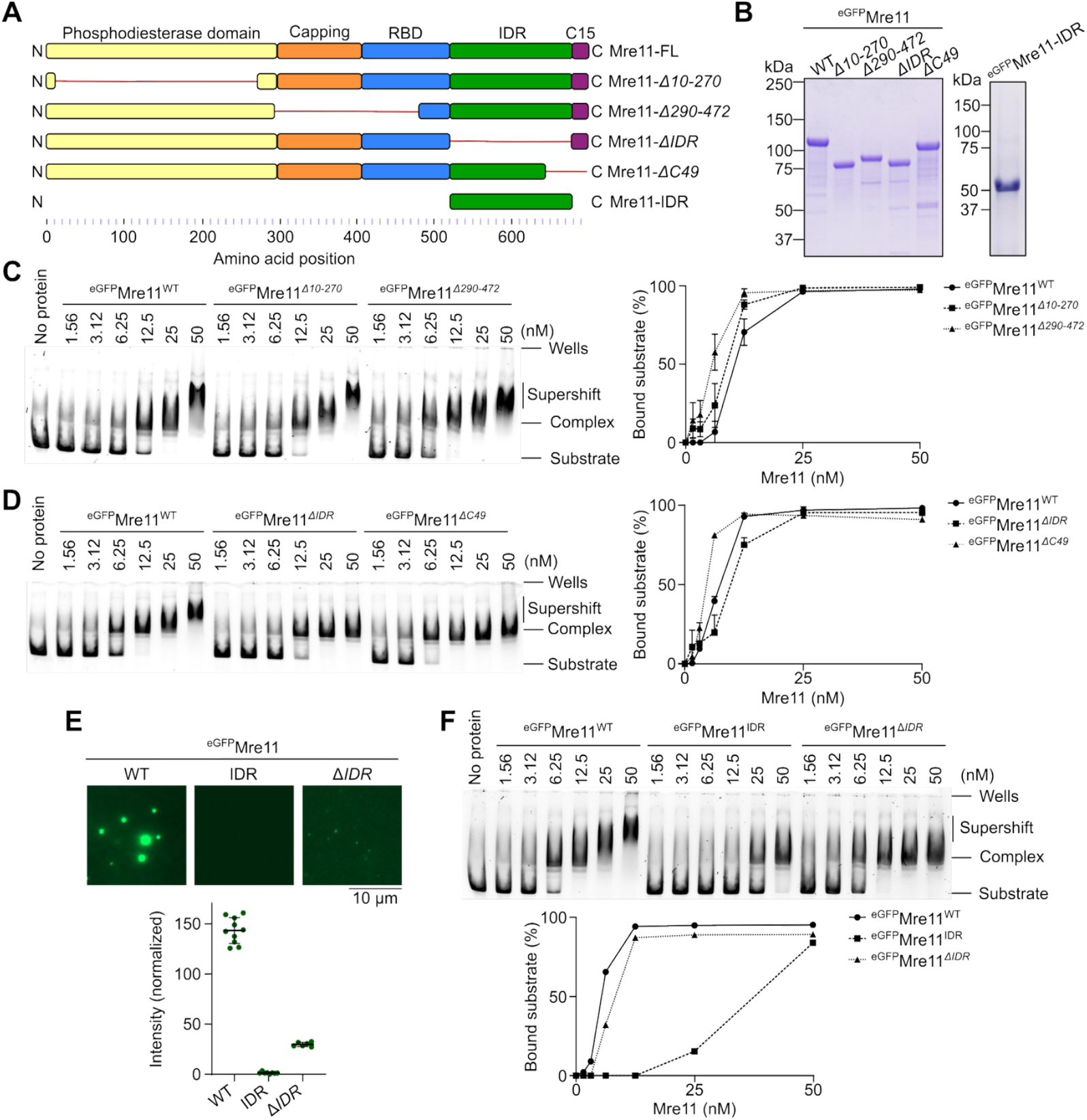
The C-terminal IDR of Mre11 is required for condensation. **(A)** Cartoon diagram of full-length and truncated Mre11. IDR ranges from residues 524-677. Red line indicates truncated regions. **(B)** SDS-PAGE of purified WT and truncated ^eGFP^Mre11. **(C, D)** Effect of Mre11 truncations on plasmid DNA binding analyzed by gel shift assay. Error bars in C and D show ranges from two independent experiments. **(E)** *In vitro* condensation analysis of eGFP-tagged Mre11, Mre11-IDR (524-677), and Mre11-*ΔIDR* (*Δ524-677*). Foci intensities are normalized to the mean of wild-type ^eGFP^Mre11. Error bars represent mean ± SD from 6-10 fields of view. **(F)** Plasmid DNA binding of eGFP-tagged Mre11-IDR in comparison with Mre11 and Mre11-*ΔIDR* analyzed by gel shift assay.

**Figure S4:**
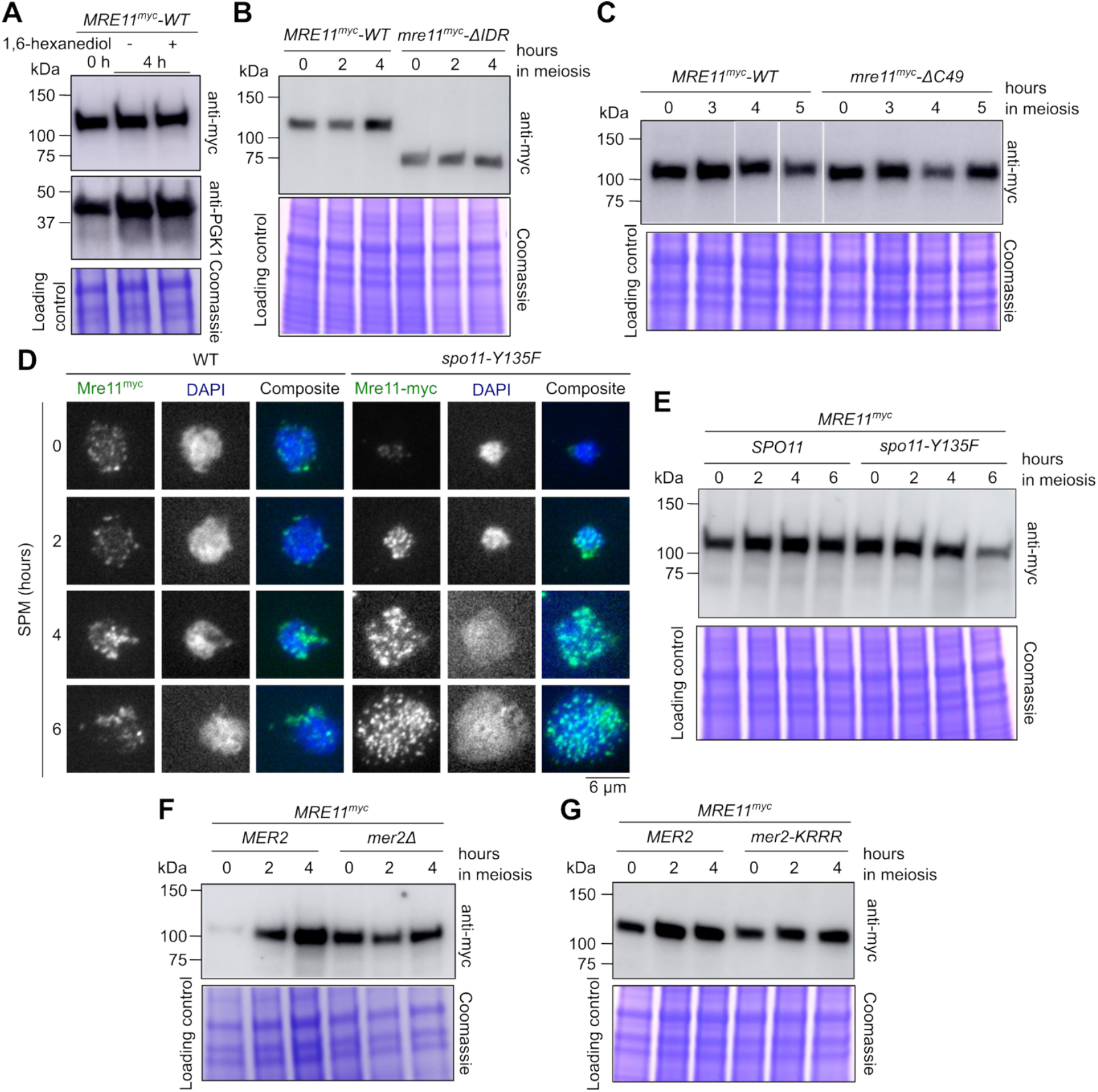
Mre11 foci formation and expression during meiosis. **(A, B, C, E, F, G)** Western blot analysis of meiotic extracts of Mre11^myc^ in (A) 1,6-hexanediol-treated Mre11^myc^ strains harvested 4 hours after transferring to SPM, (B) wild type and Mre11-*ΔIDR* strains, (C) wild type and Mre11-*ΔC49* strains, (E) wild type and *spo11-Y135F* strains, (F) wild-type and *mer2Δ* strains, and (G) wild-type and *mer2-KRRR* strains. Coomassie-stained SDS-PAGE gels and anti-PGK1 Wester blots (panel 1) serve as loading controls. In panel C, all samples were loaded on the same gel, but lanes were re-ordered. **(D)** Immunofluorescence on meiotic nuclear spreads of myc-tagged Mre11 in wild-type and *spo11-Y135F* strains. Contrary to previously published ChIP results showing similar association and dissociation kinetics of Mre11 in wild-type and *spo11-Y135F* backgrounds^40^, in our hands Mre11 foci accumulate at late time points in a *spo11-Y135F* mutant.

**Figure S5:**
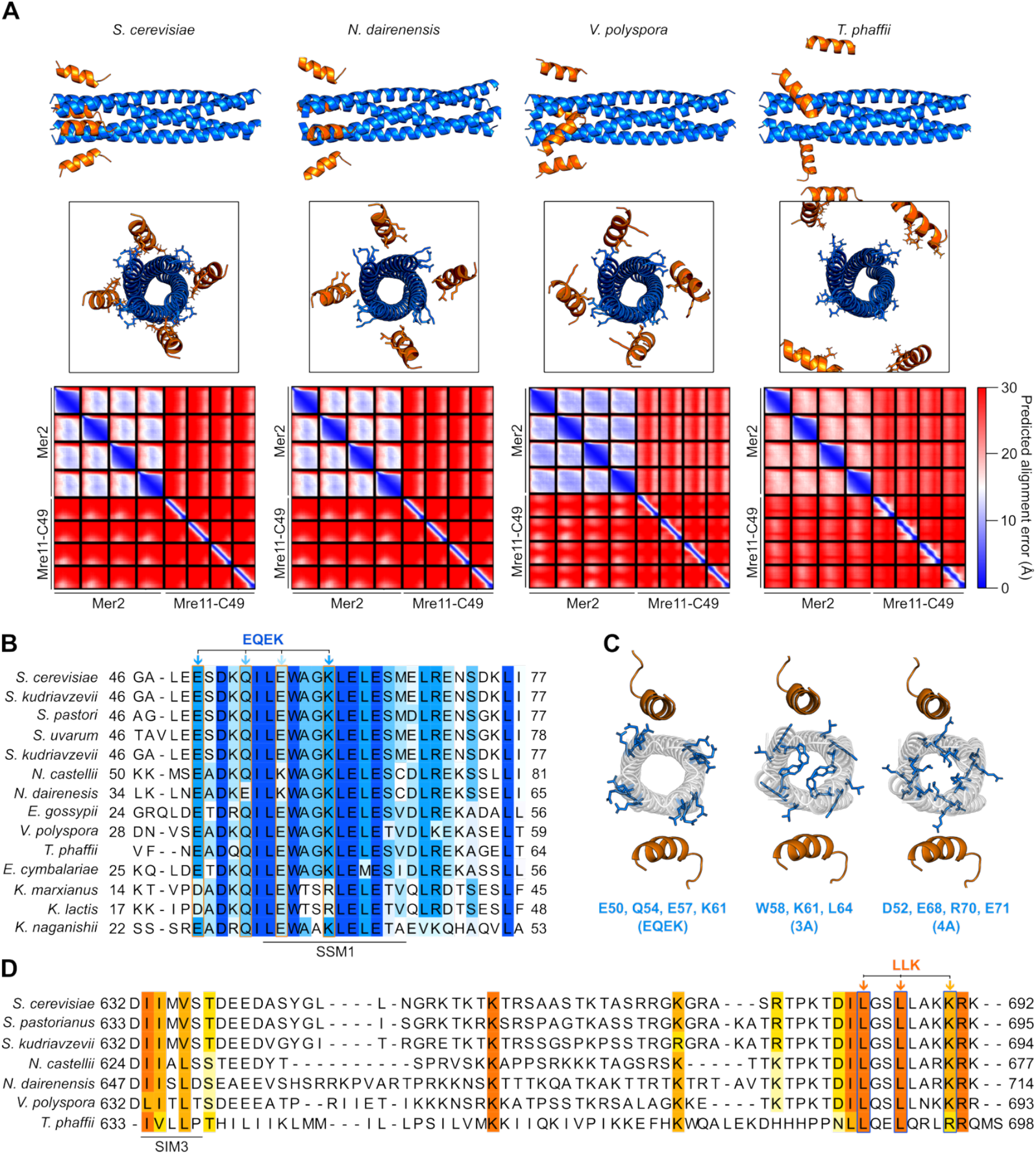
AlphaFold2 models of Mre11-Mer2 complexes and sequence conservation. **(A)** AlphaFold2 models of 4:4 Mre11-Mer2 interaction domains in various species of Saccharomycetaceae. Mre11 is shown in orange and Mer2 in blue. Mre11-LLK and Mer2-EQEK residues are shown as orange and blue sticks, respectively, in the lateral views. Disordered regions are omitted for clarity. All models are similar, except for *T. phaffi* that is aberrant. Predicted alignment error plots for each model is shown below. Dark blue represents low predicted error and high confidence, whereas lighter shades and red indicates low confidence, typical for flexible or disordered regions. NCBI accession numbers for Mer2 and Mre11, respectively, are as follows: *Saccharomyces cerevisiae* (CAA60944, BAA02017), *Naumovozyma dairenensis* (XP_003669210.1, XP_003672532.1), *Vanderwaltozyma polyspora* (XP_001647040.1, XP_001642997.1), and *Tetrapisispora phaffii* (XP_003683996.1, XP_003686402.1). The sequences used for AlphaFold modeling are provided in **Table S5. (B, D)** Multiple sequence alignments of (B) Mer2 and (D) Mre11 in members of the Saccharomycetaceae class. EQEK residues are indicated by blue arrows and orange boxes and LLK residues are indicated by orange arrows and blue boxes. Alignment is colored based on percentage identity score on Jalview with a conservation threshold of 35% for Mer2 and 50% for Mre11. The previously-identified Mer2 signature sequence motif (SSM1) is indicated^44^. **(C)** Position of Mer2 EQEK residues (blue) on the AlphaFold2 model, and comparison with published 3A and 4A mutants^41^. Mre11 is in orange and Mer2 is in white.

**Figure S6:**
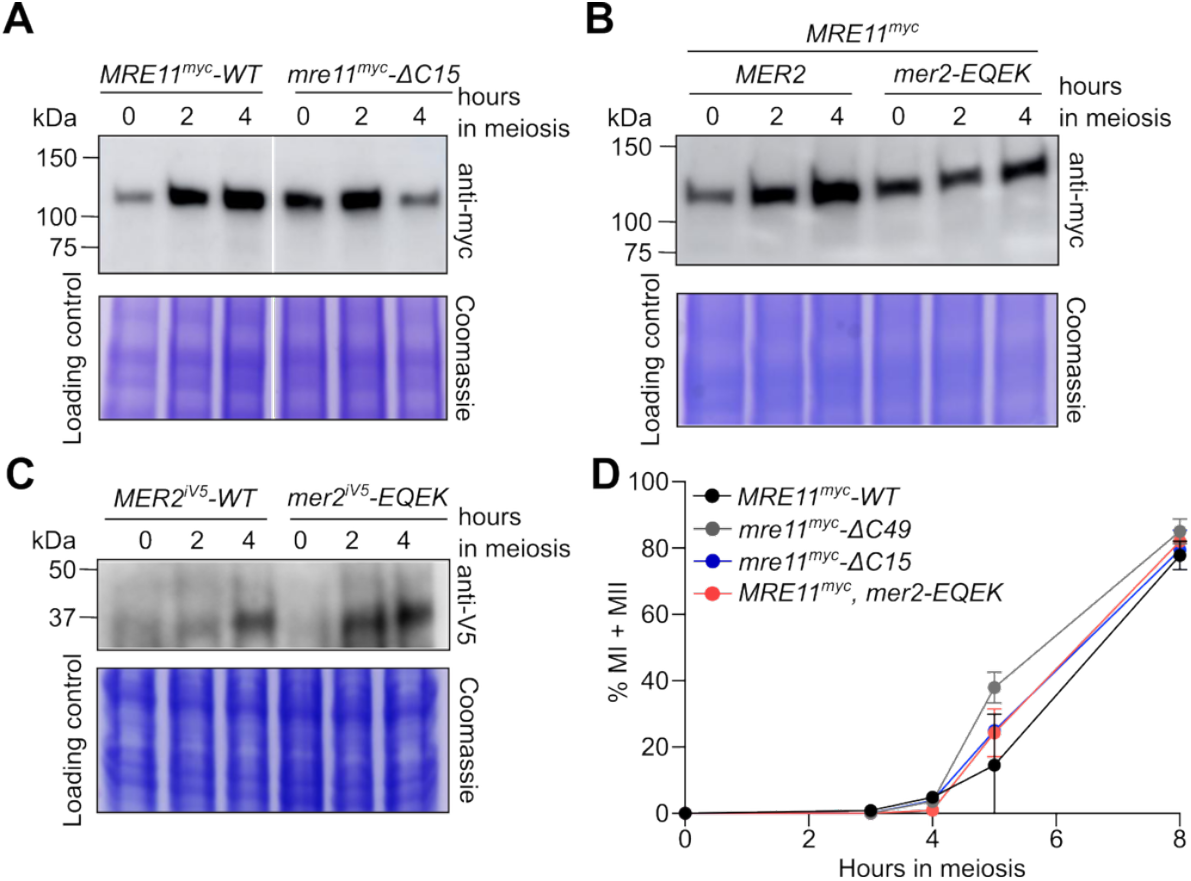
Analysis of Mer2 – Mre11 interaction mutants. **(A, B, C)** Western blot analysis of meiotic extracts of (A) *MRE11*^*myc*^*-WT* and *mre11-ΔC15*^myc^, (B) *MRE11*^*myc*^ in a *MER2* or *mer2-EQEK* strain, and (C) wild type or mutant *MER2*^*iV5*^ strains. The *MER2*^*iV5*^ allele has an internal V5 tag between Mer2 amino acids 248 and 249. Coomassie-stained SDS-PAGE gels serve as loading controls. (**D**) Meiotic progression, as indicated by the percentage of cells that have undergone the first or second meiotic divisions (MI + MII) (n = 2).

**Figure S7:**
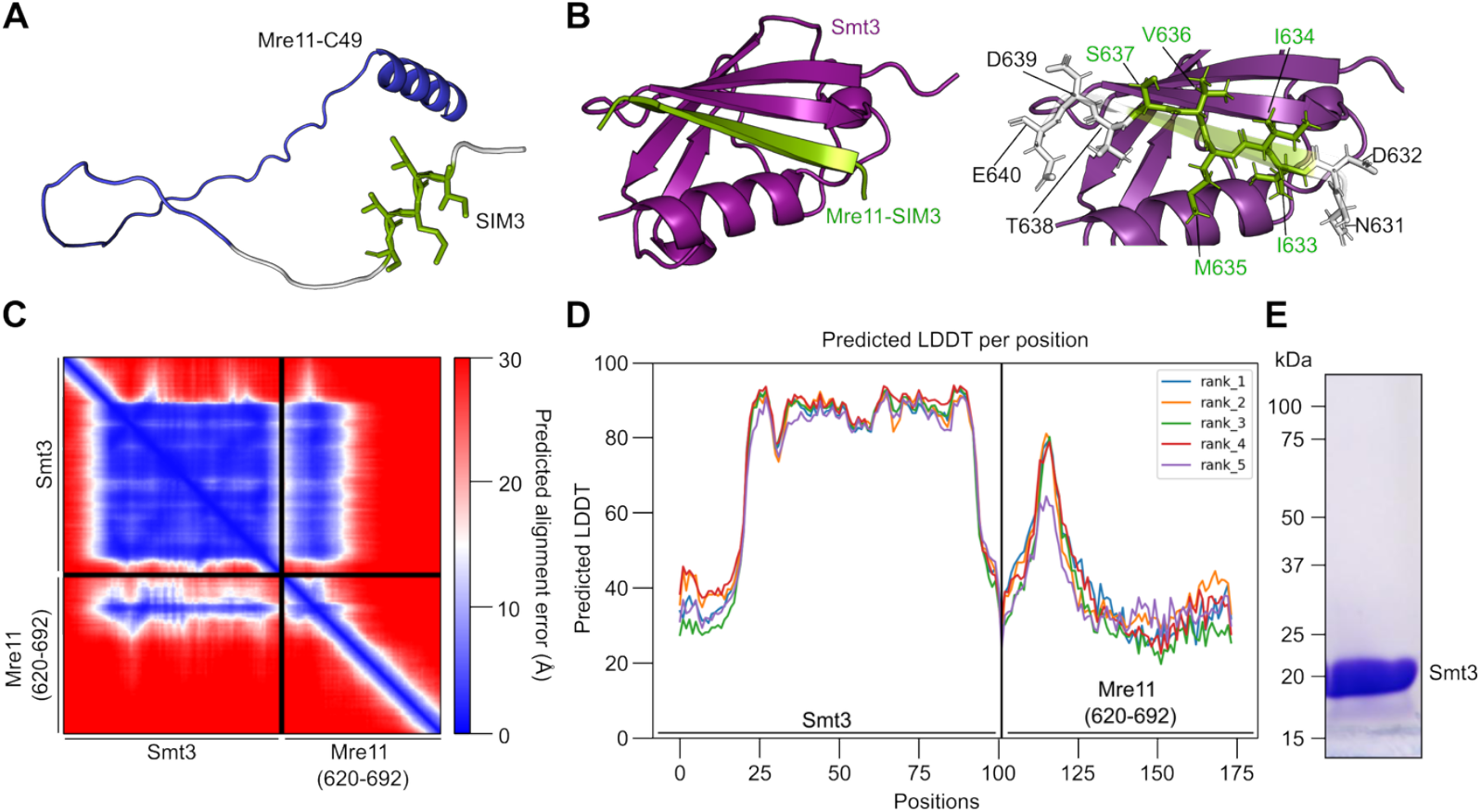
The Mre11 C-terminus contains a novel SUMO-interaction motif. **(A)** Representation of Mre11 C-terminus (residues 630-692) from AlphaFold2 database (AF-P32829-F1-v4). The SIM3 motif (green) is located immediately before the Mre11-C49 residues (blue). **(B)** AlphaFold2 model of Smt3 (purple) and Mre11-SIM3 (residues 630-640) (green). Note that SIM3 is predicted to be disordered in panel A but folds as a β-sheet when bound to Smt3. Disordered regions of Smt3 and Mre11 are omitted for clarity. The sequences used for AlphaFold modeling are provided in **Table S5. (C)** Predicted aligned error (Å) plot for AlphaFold2 model of Smt3 and Mre11-SIM3 (residues 620-692). The x- and y-axis show amino acid residue numbers. Darker blue represents low predicted error and high confidence, whereas lighter shades and red indicates low confidence, typical for flexible or disordered regions. **(D)** Predicted Local Distance Difference Test (pLDDT) plot for AlphaFold2 model of Smt3 and Mre11-SIM3. The x-axis represents amino acid position and y-axis represents per-residue pLDDT score. Higher scores indicate greater confidence in local structure prediction. **(E)** SDS-PAGE of purified His-tagged U-[^13^C,^15^N] Smt3.

**Figure S8:**
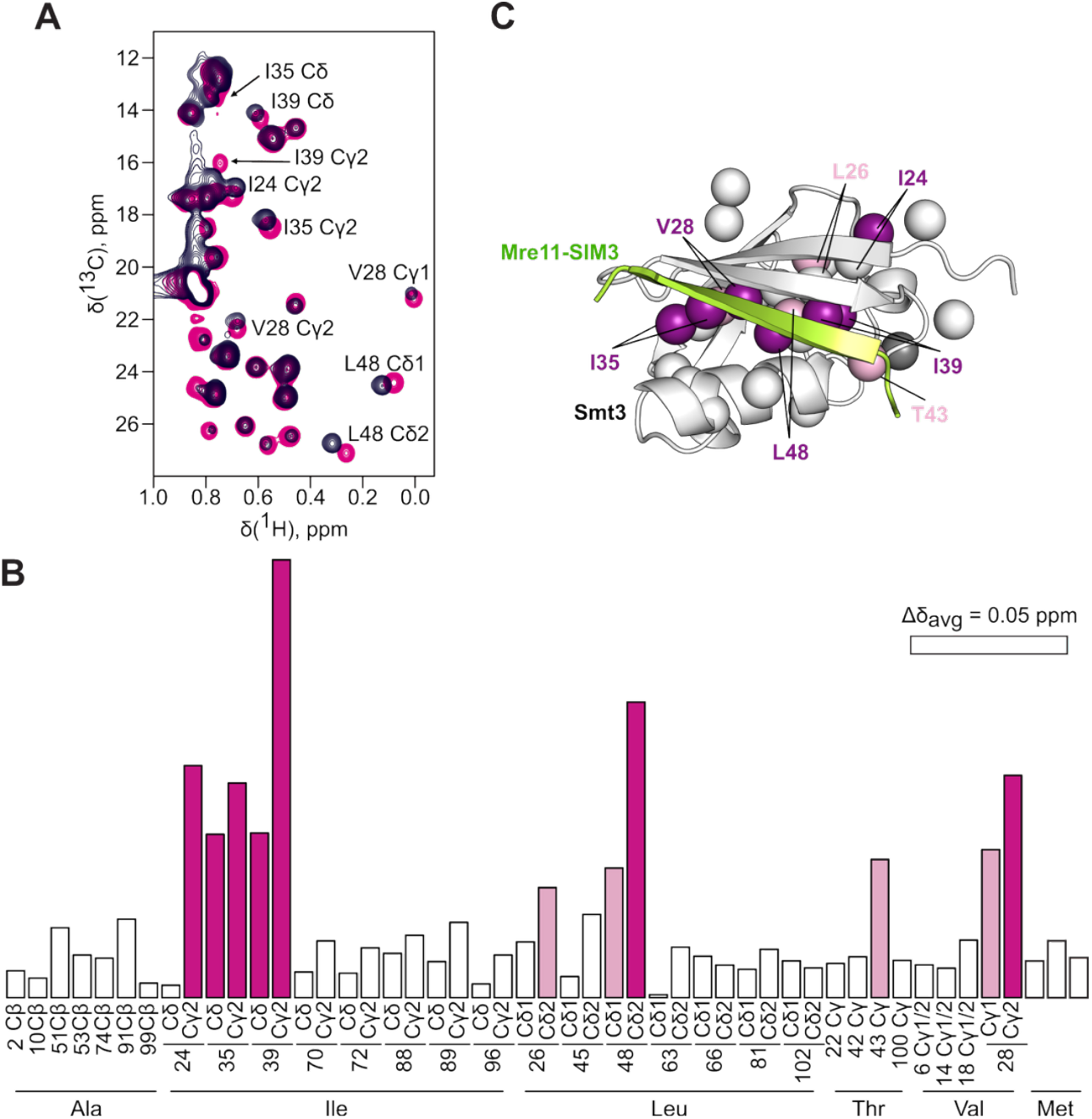
NMR analysis of Smt3 bound to wild type and mutant SIM3 peptides. **(A)** Methyl regions of [^1^H,^13^C] HSQC spectra of the free Smt3 (black) and in the presence of 1.2 molar equivalents of wild type SIM3 peptide (magenta). The labels indicate the protein CH_3_ groups showing the largest binding shifts. **(B)** Average methyl chemical shift perturbations (Δδ_avg_) of Smt3 upon binding to wild-type SIM3 peptide. The pink and magenta bars correspond to the CH_3_ groups with Δδ_avg_ > 0.03 ppm and > 0.05 ppm, respectively. For V6, V14, and V18, which show identical Cy1 and Cy2 NMR resonances, a single bar is shown. As the Cδ resonances of M49, M60, and M83 were not explicitly assigned in this work, the data for the methionine methyls are represented by unmarked bars. **(C)** Chemical shift mapping of the wild type SIM3 peptide binding. Smt3 methyls are shown as spheres, colored according to Δδ_avg_ as in panel B. The bound SIM3 peptide is in green, and the disordered Smt3 N- and C-termini are omitted for clarity.

**Figure S9:**
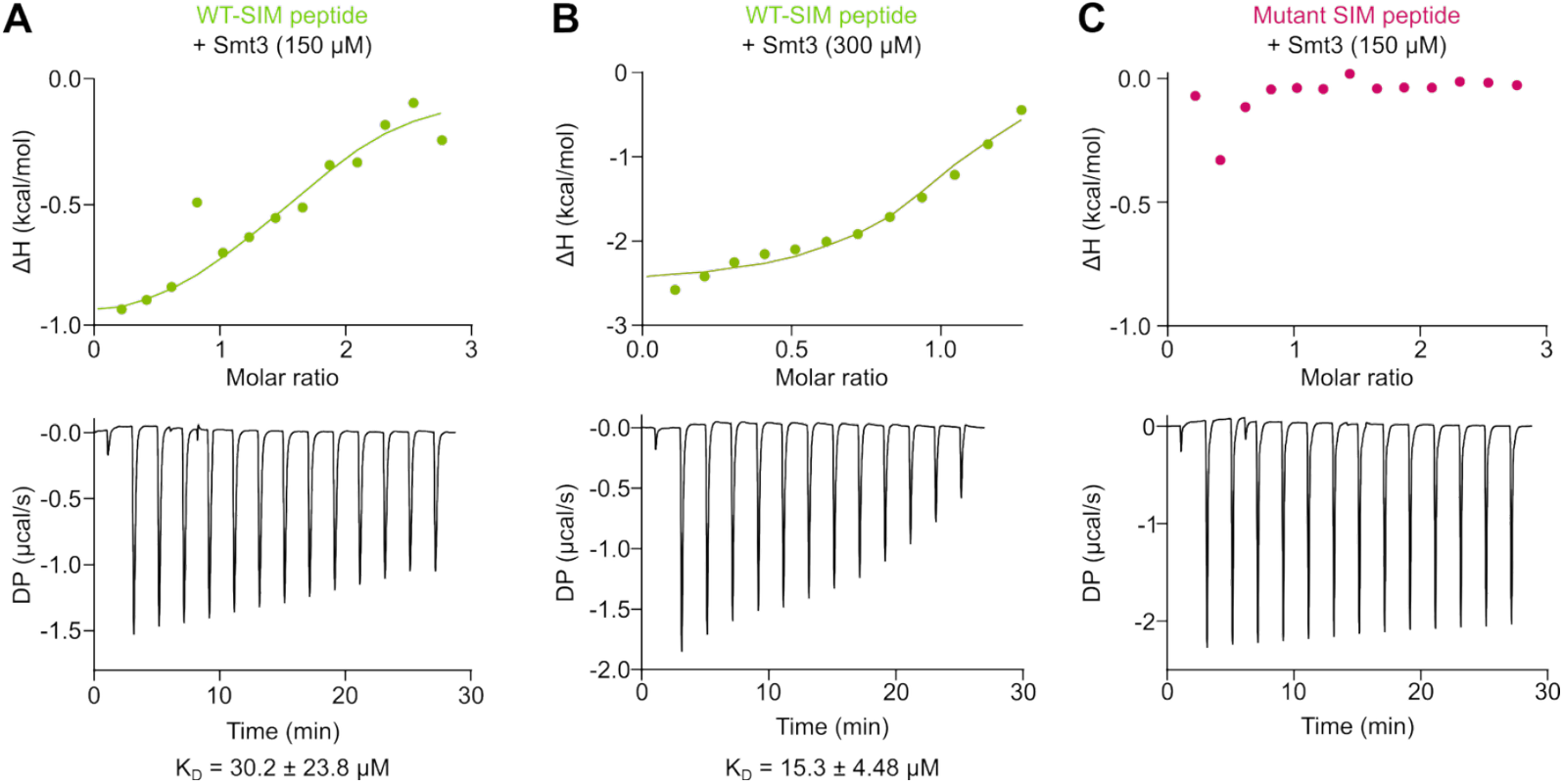
ITC analysis of Smt3 binding to wild-type or mutant SIM3 peptides. Top panels, integrated heat peaks ΔH (kcal/mol) as a function of molar ratio (peptide/protein concentration) after buffer subtraction and offset correction. Bottom panels, raw data plots indicating differential power (μcal/s) after baseline correction in function of time. The data was fitted to a single binding site model and the number of binding sites (N) was set to 1. The model allowed for the calculation of the equilibrium dissociation constant K_D_ as provided below each graph.

**Figure S10:**
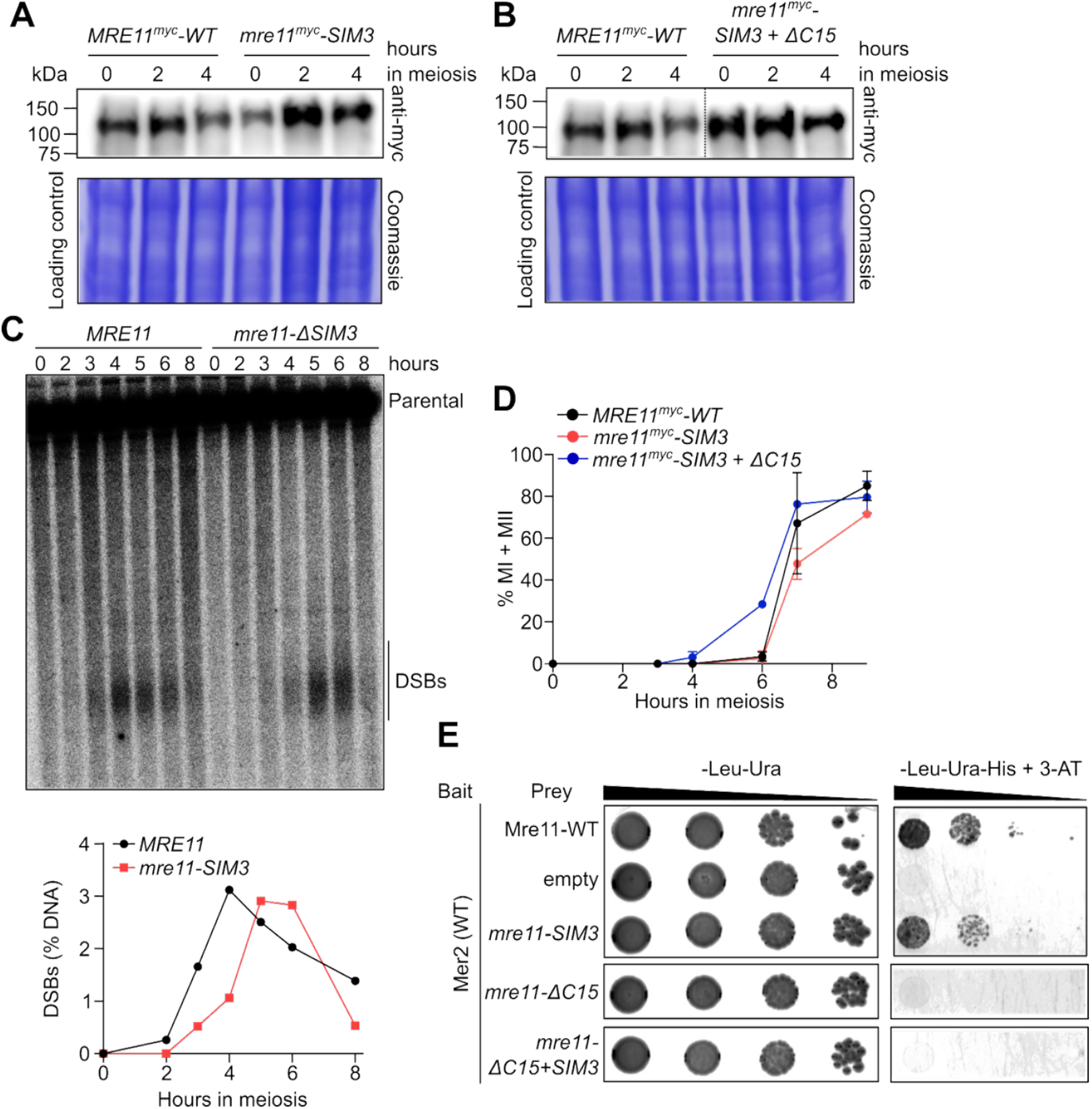
Protein expression and meiotic progression of Mre11 SIM3 mutants. **(A, B)** Western blot analysis of meiotic extracts of (A) *MRE11*^*myc*^*-WT* and *mre11-SIM3*^myc^, (B) *MRE11*^*myc*^*-WT* and *mre11-SIM3+C15*^myc^. Coomassie-stained SDS-PAGE gels serve as loading controls. **(C)** Southern blot analysis of meiotic DSB formation at the *GAT1* hotspot. **(D)** Meiotic progression, as indicated by the percentage of cells that have undergone the first or second meiotic divisions (MI + MII) (n = 2). (**E**) Yeast-two-hybrid analysis between Mer2 wild-type (binding domain, bait) and Mre11 wild-type and mutants (activation domain, prey).

**Table S1.**
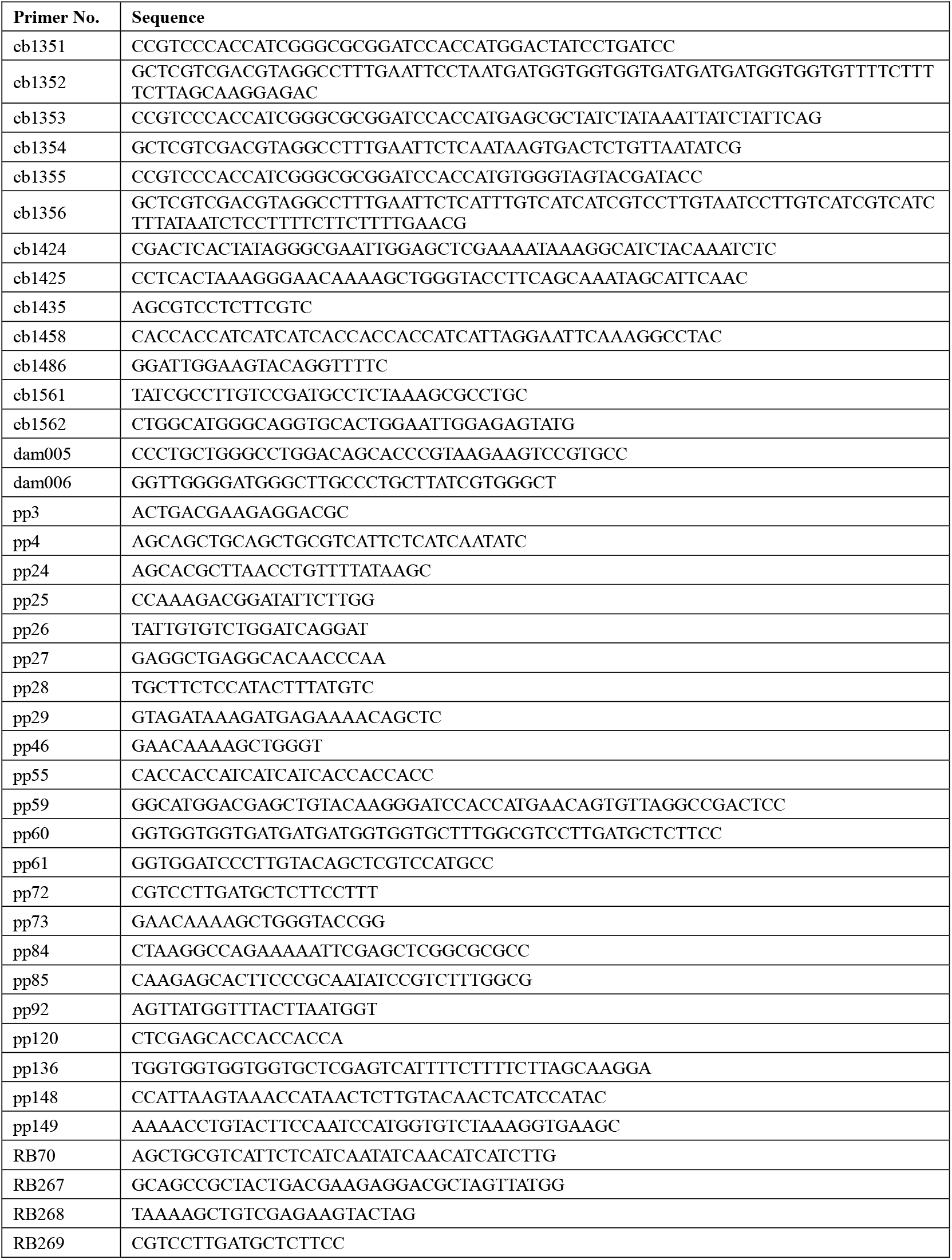
Oligonucleotides used in this study.

**Table S2.**
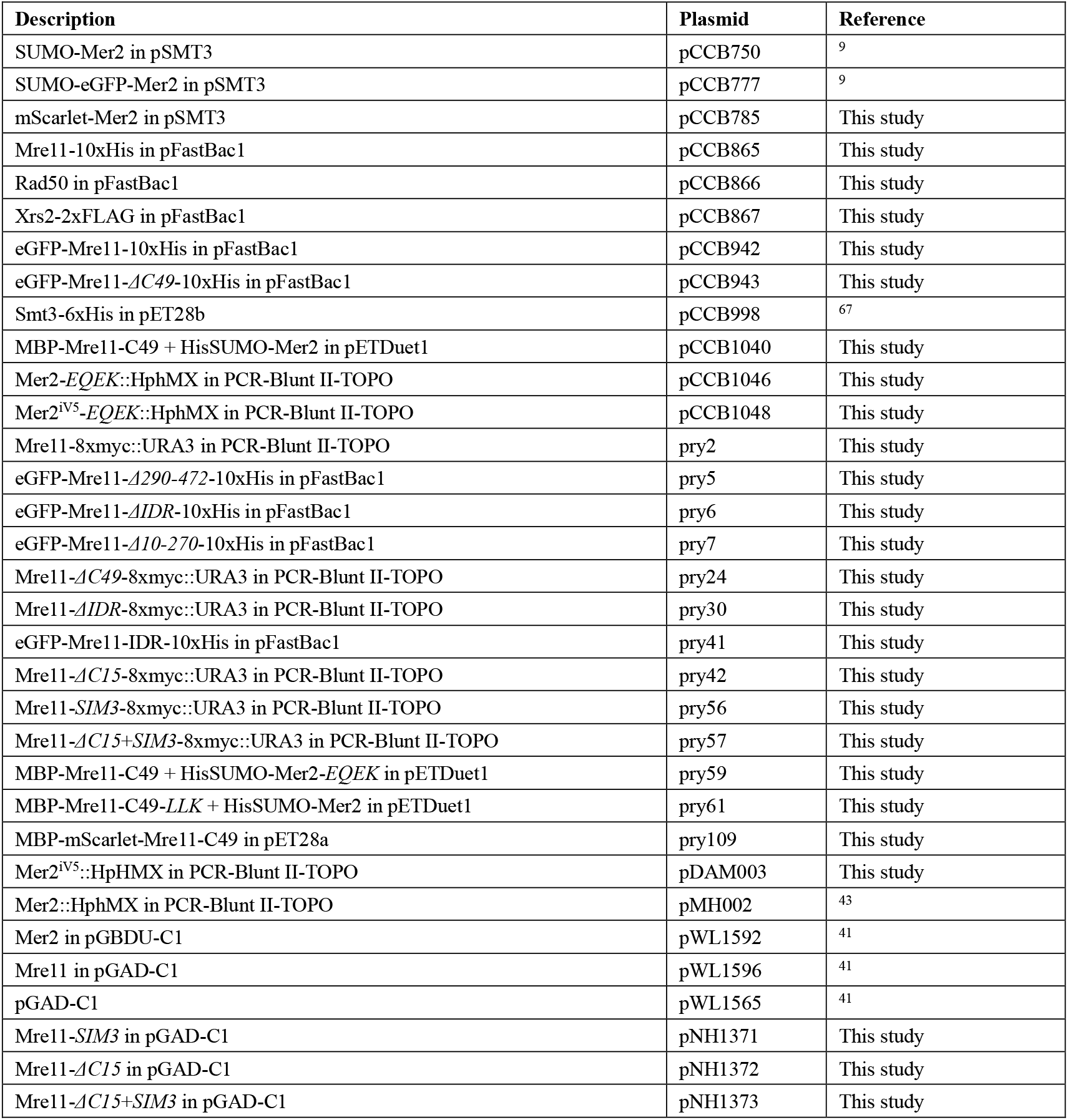
Plasmids used in this study.

| Description | Plasmid | Reference |
| --- | --- | --- |
| SUMO-Mer2 in pSMT3 | pCCB750 | <sup>9</sup> |
| SUMO-eGFP-Mer2 in pSMT3 | pCCB777 | <sup>9</sup> |
| mScarlet-Mer2 in pSMT3 | pCCB785 | This study |
| Mre11-10xHis in pFastBac1 | pCCB865 | This study |
| Rad50 in pFastBac1 | pCCB866 | This study |
| Xrs2-2xFLAG in pFastBac1 | pCCB867 | This study |
| eGFP-Mre11-10xHis in pFastBac1 | pCCB942 | This study |
| eGFP-Mre11-*ΔC49*-10xHis in pFastBac1 | pCCB943 | This study |
| Smt3-6xHis in pET28b | pCCB998 | <sup>67</sup> |
| MBP-Mre11-C49 + HisSUMO-Mer2 in pETDuet1 | pCCB1040 | This study |
| Mer2-*EQEK*::HphMX in PCR-Blunt II-TOPO | pCCB1046 | This study |
| Mer2<sup>iV5</sup>-*EQEK*::HphMX in PCR-Blunt II-TOPO | pCCB1048 | This study |
| Mre11-8xmyc::URA3 in PCR-Blunt II-TOPO | pry2 | This study |
| eGFP-Mre11-*Δ290-472*-10xHis in pFastBac1 | pry5 | This study |
| eGFP-Mre11-*ΔIDR*-10xHis in pFastBac1 | pry6 | This study |
| eGFP-Mre11-*Δ10-270*-10xHis in pFastBac1 | pry7 | This study |
| Mre11-*ΔC49*-8xmyc::URA3 in PCR-Blunt II-TOPO | pry24 | This study |
| Mre11-*ΔIDR*-8xmyc::URA3 in PCR-Blunt II-TOPO | pry30 | This study |
| eGFP-Mre11-IDR-10xHis in pFastBac1 | pry41 | This study |
| Mre11-*ΔC15*-8xmyc::URA3 in PCR-Blunt II-TOPO | pry42 | This study |
| Mre11-*SIM3*-8xmyc::URA3 in PCR-Blunt II-TOPO | pry56 | This study |
| Mre11-*ΔC15*+*SIM3*-8xmyc::URA3 in PCR-Blunt II-TOPO | pry57 | This study |
| MBP-Mre11-C49 + HisSUMO-Mer2-*EQEK* in pETDuet1 | pry59 | This study |
| MBP-Mre11-C49-*LLK* + HisSUMO-Mer2 in pETDuet1 | pry61 | This study |
| MBP-mScarlet-Mre11-C49 in pET28a | pry109 | This study |
| Mer2<sup>iV5</sup>::HpHMX in PCR-Blunt II-TOPO | pDAM003 | This study |
| Mer2::HphMX in PCR-Blunt II-TOPO | pMH002 | <sup>43</sup> |
| Mer2 in pGBDU-C1 | pWL1592 | <sup>41</sup> |
| Mre11 in pGAD-C1 | pWL1596 | <sup>41</sup> |
| pGAD-C1 | pWL1565 | <sup>41</sup> |
| Mre11-*SIM3* in pGAD-C1 | pNH1371 | This study |
| Mre11-*ΔC15* in pGAD-C1 | pNH1372 | This study |
| Mre11-*ΔC15*+*SIM3* in pGAD-C1 | pNH1373 | This study |

**Table S3.**
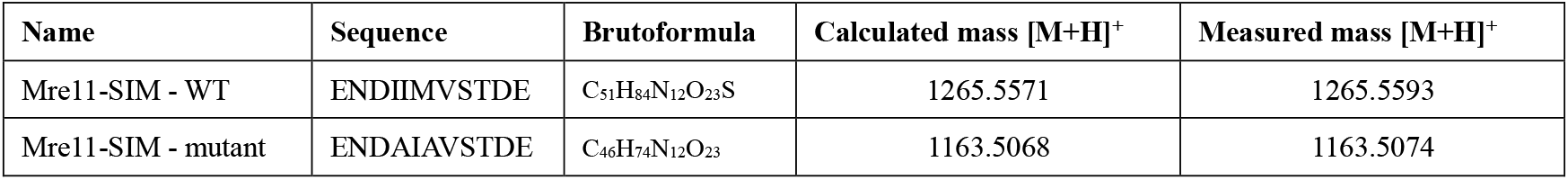
Synthetic peptides used in this study.

**Table S4.**
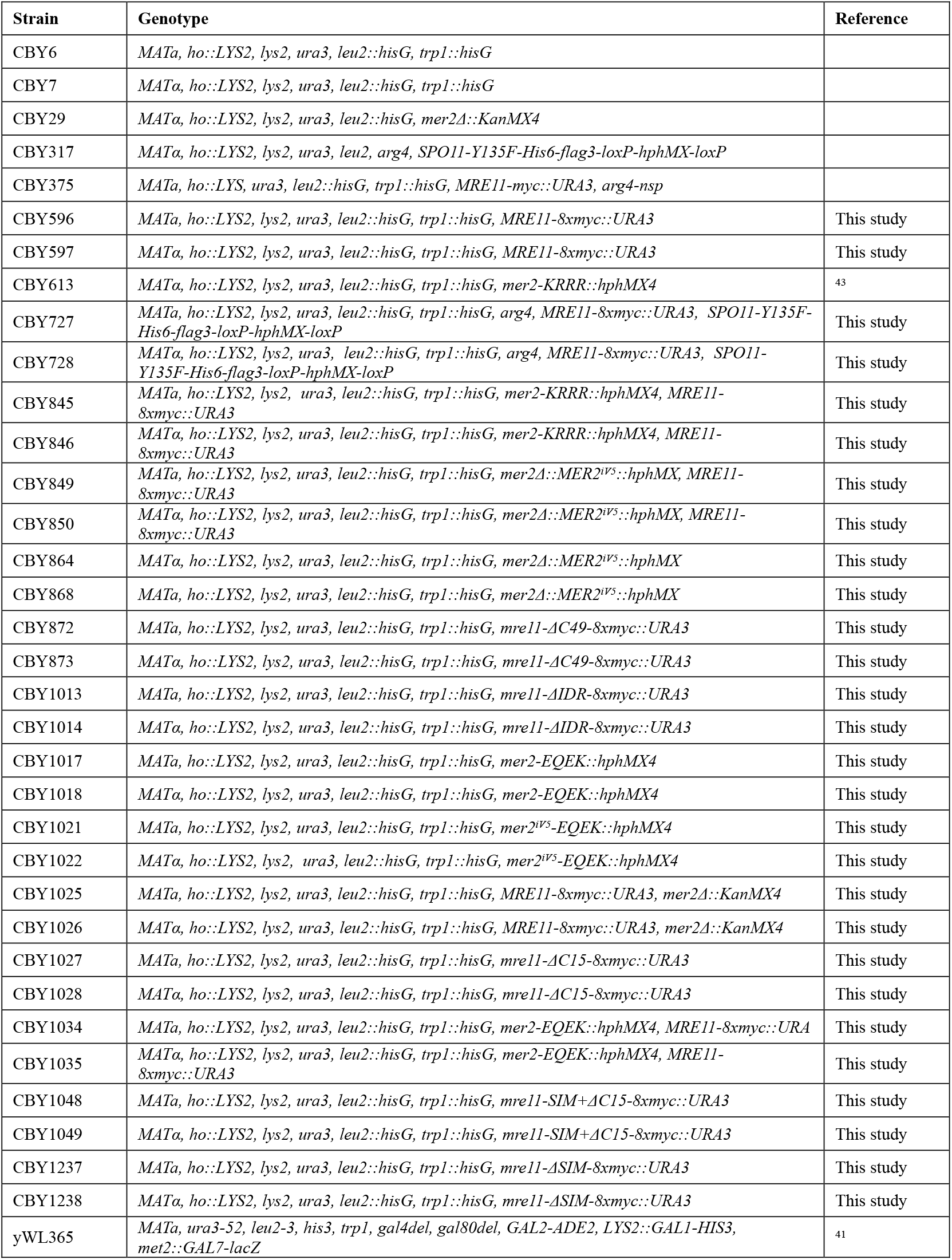
Yeast strains used in this study.

**Table S5.**
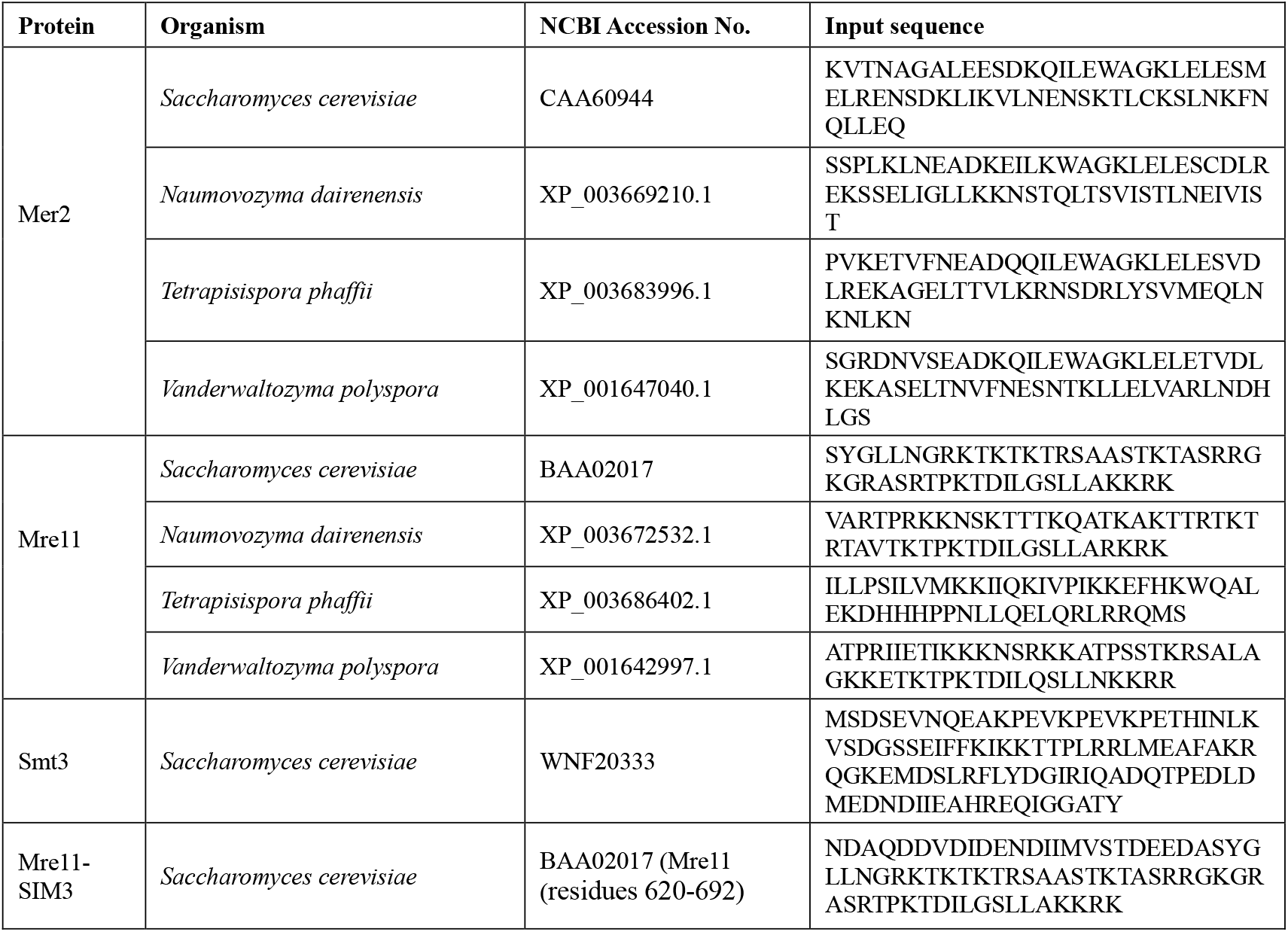
Protein sequences used for AlphaFold modeling.

## Notes

### Competing Interest Statement

The authors have declared no competing interest.

